# Distinct virus-derived circular RNA molecule influences host response during SARS-CoV-2 infection

**DOI:** 10.64898/2026.04.28.721328

**Authors:** Elysse N. Grossi-Soyster, Rebekah C. Gullberg, Arjun Rustagi, Jae Seung Lee, Catherine A. Blish, Sara Cherry, Julia Salzman, Peter Sarnow

**Author notes:** Corresponding author: Dr. Peter Sarnow, Department of Microbiology and Immunology, Stanford University School of Medicine, Stanford, CA 94305, USA. Department of Medicine, University of California, San Francisco, San Francisco, CA 94121. Center for Infectious Diseases Vaccine and Diagnosis Innovation (CEVI), Korea, Research Institute of Chemical Technology (KRICT), Daejeon, 34114, Republic of Korea.

## Abstract

Virus-derived circular RNA molecules (VcircRNAs) are expressed by many RNA viruses during infection. Putative functions include modulating viral replication and interacting with the host immune response. Some function as non-coding RNA fragments that regulate gene expression through binding to complementary RNA sequences, whereas others contain internal ribosomal entry site (IRES) sequences or non-canonical modifications that allow them to be translated. Here, we confirm the expression of a distinct SARS-CoV-2 VcircRNA molecule, circ7b8N, that has not been previously identified. We found that circ7b8N is expressed and detectable in cell culture infections and in acute infections across SARS-CoV-2 variants and shows promise for detection in post-acute clinical samples. Conservation of circ7b8N junctions is limited to the nearest phylogenetic relatives within the betacoronavirus genus but are present in other human and bat-infecting coronaviruses. Host cell gene expression is modulated by the treatment with circ7b8N agnostic of viral infection. The discovery and subsequent confirmation of circ7b8N expressed by SARS-CoV-2 provides a new biomarker for infection, and its conservation across variants suggests functional importance.

**Author Summary:** Circular RNAs are a well-documented class of molecules expressed by mammalian cells. However, circular RNA molecules expressed by RNA viruses remain largely uncharacterized regarding their generation, specific functions, and roles in host-pathogen interactions. Our computational predictions discovered thousands of distinct circular RNA molecules expressed by SARS-CoV-2. Among these, we confirmed the presence of circ7b8N, the most abundant SARS-CoV-2-derived circular RNA identified in our sequencing data. We found that circ7b8N localizes outside the nucleus and is detectable in clinical samples collected both during and after acute SARS-CoV-2 infection. Although overexpression of circ7b8N was not found to alter viral titers, it modulated the expression of host genes related to immune response activation and membrane remodeling. This suggests that circ7b8N may simultaneously provide pro- and anti-viral functions independent of influencing viral replication. Phylogenetic analyses of coronaviruses suggest that the expression of circ7b8N is a relatively recent evolutionary event, and it is conserved across SARS-CoV-2 variants from the first five years of the pandemic. The abundant presence of circ7b8N across variants in both sequencing data and clinical samples implies it plays a multifaceted role in SARS-CoV-2 pathogenesis.

## Introduction

Since emerging in 2019[1], Severe Acute Respiratory Syndrome Coronavirus 2 (SARS-CoV-2) has exemplified rapid evolution and viral persistence resulting in continued global spread. SARS-CoV-2 is a positive-sense, single-stranded RNA virus in the *Nidoviridae* order, *Coronaviridae* family which utilizes double membrane vesicles to replicate in the cytoplasm of mammalian host cells[2–4]. Despite the extensive, collaborative scientific and public health responses that supported the rapid development of SARS-CoV-2 vaccines and therapeutics, much of the RNA landscape that shapes SARS-CoV-2 infection dynamics remains elusive.

Circular RNA molecules (circRNA), typically generated during host pre-mRNA splicing by a backsplicing mechanism[5], are also expressed from many viral DNA genomes[6]. The discovery of VcircRNAs has added a new component to previously characterized viral lifecycles and host-virus interactions. Also, many positive-stranded RNA viruses, including hepatitis C virus (HCV)[7], appear to generate hundreds of VcircRNAs from fragments of their viral genome, likely by a novel cytoplasmic splicing mechanism. Mechanisms of VcircRNA biogenesis have yet to be determined but likely involve RNA structures and distinct RNA nucleases and ligases. Thus, it is likely that many RNA viruses have adapted niche functions to support specific immune evasion and replication processes. Early evidence suggests that some circRNA and VcircRNA functions include acting as a sponge for microRNAs[8] and ribosomal binding proteins, regulating transcription and translation[5,9–12]. While VcircRNA research is in its nascent stages, previous studies emphasize computational prediction of VcircRNAs and allude to potential functions based on RNA binding protein motifs. Previous work by Cao et al. indicates VcircRNAs expressed by HCV enhance viral replication, and specific HCV VcircRNAs containing an internal ribosomal entry site (IRES) can be translated into unique viral proteins[7]. Beyond directly impacting viral replication, VcircRNAs have been overlooked as potential biomarkers for infection, as has been proposed for host circRNAs in monitoring overall health and disease[13,14]. Others have predicted VcircRNAs expressed by coronaviruses (CoVs), yet no attempts to detect and verify VcircRNAs from SARS-CoV-2 from acute infections have been attempted previously[15,16]. Therefore, we hypothesize that (1) SARS-CoV-2 expresses VcircRNAs, and that (2) such VcircRNAs modulate SARS-CoV-2 infections, thus possibly driving epidemiological trends in global transmission. With this work, we confirmed the expression of a distinct SARS-CoV-2 VcircRNA, circ7b8N, in cell culture and clinical isolates, and identified the impacts of circ7b8N on viral kinetics and variant persistence.

## Results

### Prediction of SARS-CoV-2 VcircRNAs

To predict potential VcircRNAs expressed by SARS-CoV-2, we infected A549-hACE2 cells with the WA1 strain of SARS-CoV-2, harvested total RNA for sequencing (RNAseq), and processed the RNAseq data using reference-dependent computational algorithms (SICILIAN[17], CIRI2[18,19], circRNA_finder[20], and find_circ[21]) specifically designed for detecting circRNAs and alternative splicing products. Initial predictions for SARS-CoV-2 were processed using SICILIAN after alignment with the primary reference genome Hu.1 from 2019 (GenBank #NC_045512.2). SICILIAN generated a list of thousands of positive and negative SARS-CoV-2 VcircRNAs, 2,060 and 3,264 respectively, in varying sequence lengths (Fig. 1). We then used the VcircRNA prediction data to investigate potential trends associated with or pre-empting circularization, whether in sequence length (Fig. 1A), nucleotide sequence, or region of the genome from which such VcircRNAs were predicted. We noticed a high level of diversity across all conditions, as VcircRNAs were predicted with near complete coverage of the entire SARS-CoV-2 genome (Fig. 1B), and with varying sequence lengths (Fig. 1A-B). VcircRNAs are predicted in every ORF, yet a concentration is found in the structural proteins and accessory factors region in the 3’ end of the genome, with heavy coverage in the nucleocapsid (N) ORF (Fig. 1B). Large arcs spanning almost the entirety of the genome (5’ - 3’) are likely products of discontinuous replication for the generation of subgenomic RNAs (sgRNAs) rather than VcircRNAs encompassing nearly the entire genome.

**Figure 1:**
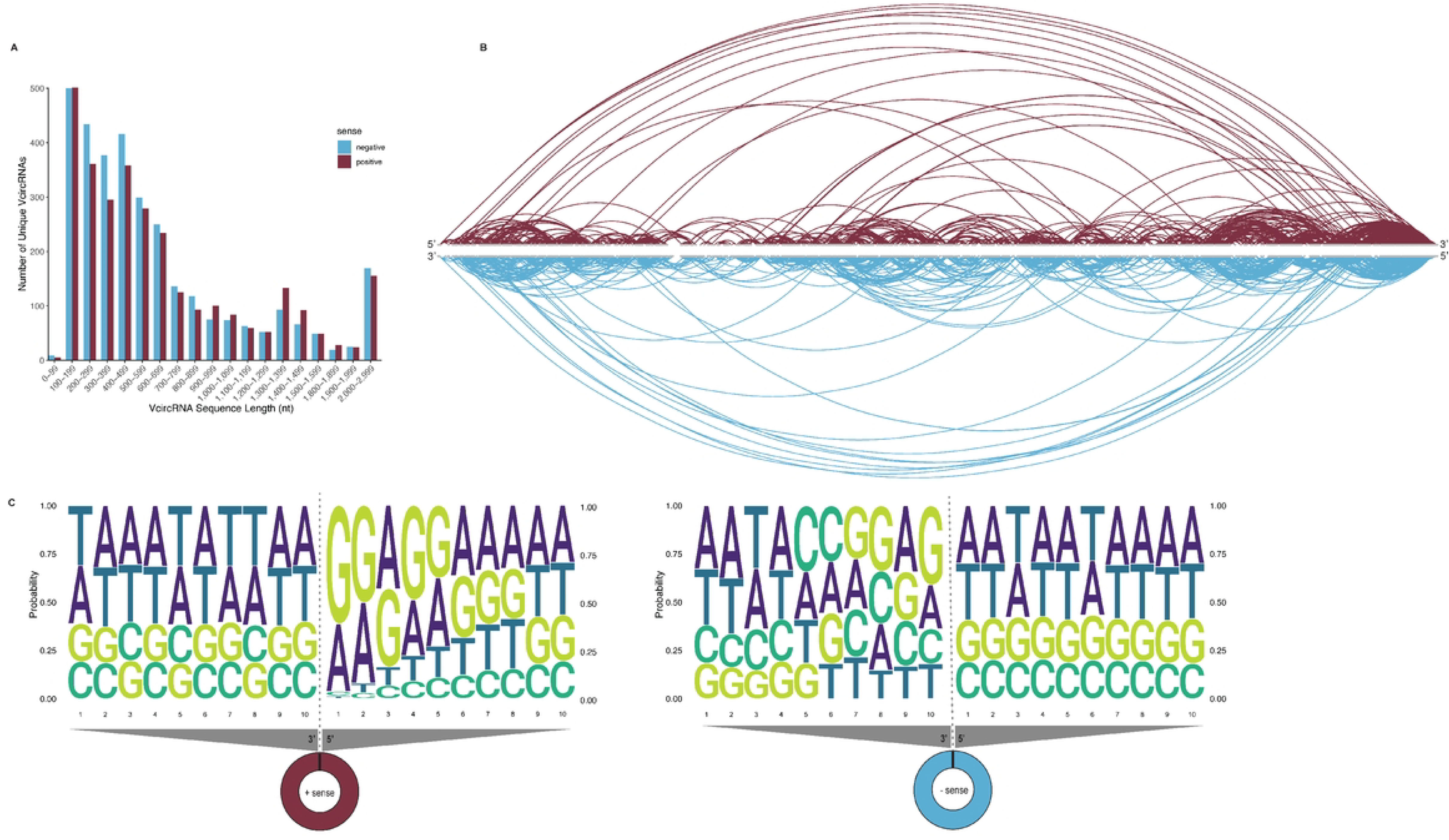
SICILIAN predictions of VcircRNAs from SARS-CoV-2. (A) Reported lengths of predicted SARS-CoV-2 VcircRNAs. Each putative VcircRNA, as predicted by SICILIAN, were comprised of variable nucleotide lengths shown on the x-axis. Positive-sense VcircRNAs (maroon) were primarily 100 - 699 nucleotides in length, or 1,300 - 1,399 nucleotides in length. Negative-sense VcircRNAs (light blue) were mostly shorter lengths, ranging from 100-699 nucleotides, with a majority skewing towards the 100-299 nucleotide range. (B) Arc diagram of predicted VcircRNAs across the SARS-CoV-2 genome. VcircRNAs are predicted across the SARS-CoV-2 genome in both positive- and negative-sense orientations, shown in maroon (upper arcs) and light blue (lower arcs), respectively. (C) Sequence logos were constructed using RNAseq data to analyze nucleotide motifs of SICILIAN-predicted SARS-CoV-2 VcircRNA junction sites. Probabilities were calculated for each nucleotide within 10 positions from the terminal end of each breakpoint. Thymine (T) is reported here in Uracil (U) positions because of RNAseq processing.

Prediction capabilities of other reference-dependent algorithms CIRI2, circRNA_finder, and find_circ were compared to initial SICILIAN predictions, and further compared to the predictions by Cai et al.[15] that were based on Calu3 cells infected with SARS-CoV-2 (Fig. S1). Each algorithm predicted circRNAs generated across the SARS-CoV-2 genome, but very few predicted VcircRNAs were predicted by multiple algorithms (Fig. S2). CIRI2 predicted 72 positive sense and 176 negative sense VcircRNAs (Fig. S1A), some of which had paired breakpoints present on the complementary strands as seen in SICILIAN predictions. Many of the negative sense VcircRNAs predicted by CIRI2 pair 5’ breakpoints in ORF1ab with 3’ breakpoints in the accessories and structural proteins region of the genome, predicting large VcircRNAs representing larger spans of the genome sequence.

Algorithm circRNA_finder heavily favored prediction of positive sense VcircRNAs, with 1,751 positive sense and only 2 negative sense VcircRNAs predicted (Fig. S1C), suggesting bias towards positive strand detection or restrictive specificity for negative sense predictions. Both negative sense VcircRNAs predicted by circRNA_finder are located in the 3’ region of the genome, one spanning 400 nucleotides from ORF M to ORF 6, and the other comprised of roughly 250 nucleotides in ORF N. CircRNA_finder was not only prolific at predicting VcircRNAs from the positive sense strand of the genome, but also at predicting VcircRNAs that were much longer in nucleotide sequence length. Compared to other algorithms, circRNA_finder predicted many larger VcircRNAs that bridged thousands of nucleotides across the genome. Prediction by find_circ indicated 304 and 796 VcircRNAs from positive sense and negative sense strands, respectively, many of which with complementary breakpoints suggesting positive and negative sense complements could be produced of the same VcircRNA (Fig. S1E).

To identify whether there are sequence motifs that are more likely to comprise the 5’ or 3’ breakpoints or the complete contiguous junction sites, probabilities were calculated for the ten nucleotides on either end of the junction site, specifically the ten terminal nucleotides the 5’ and 3’ ends of the junction breakpoints (Figs. 1C and S1). For the SICILIAN predicted positive-sense VcircRNAs (Fig. 1C), probabilities across adenosine (A), guanine (G), cytosine (C), and uracil (U) (shown as thymine (T) due to RNAseq preparation) were roughly equal at the 3’ breakpoint nucleotide sequence, indicating a random chance that any of the four nucleotides could be placed along the 3’-most nucleotides, and that there is no specific motif or pattern to indicate a breakpoint. However, on the 5’ end of the positive-sense VcircRNAs, the ten most 5’ terminal nucleotides heavily favored a string of guanine in the first, second and fourth nucleotide positions (60%, 50%, and 50% probabilities, respectively), followed by adenosine selection in the first, second, and third positions (35%, 40%, and 45% probabilities, respectively). SICILIAN predicted minus-sense VcircRNAs (Fig. 1C), the nucleotide immediately flanking the 3’ breakpoint was found to have a 40% probability for guanine, yet beyond that nucleotide position, significant, identifiable motifs at either 3’ or 5’ nucleotide were not observed. Upon RNA binding protein motif analysis, the probable GGAA motif of the positive-sense 5’-end nucleotides is a match for eukaryotic translation initiation factor 4B (EIF4B), an RNA binding protein that specializes in translation initiation.

Other algorithms generally did not display any notable motifs along the terminal ten nucleotides at each breakpoint (Fig. S1) with the exception of CIRI2, which reported more predicted VcircRNAs with terminal guanine in the positive sense 3’ breakpoint (50%), the positive sense 5’ breakpoint (60%), and the negative sense 3’ breakpoint (75%) (Fig. S1B). Upstream from the terminal nucleotide, all three algorithms illustrated uracil (U, shown as thymine (T)) repeats in the nucleotide positions 2 to 4, 5, or 6 in the 5’ breakpoints, and in nucleotides 5 through 10 for the positive sense 3’ ends predicted by find_circ. These UUU repeats may provide a binding motif for the La proteins that are known to chaperone and shuttle RNA molecules between the cytoplasm and the nucleus[22]. The rest of this work concentrates on the VcircRNA predicted by SICILIAN.

### Confirmation of circ7b8N

Quantification of predicted VcircRNA abundance was captured by RNAseq read counts, indicating changes in expression across infection timepoints (24-, 48-, and 72-hours post infection (hpi)) (Fig. 2). Read counts ranged from 2 to 560 reads for positive strand VcircRNAs, and 2 to 190 reads for negative strand VcircRNAs. Since circRNAs are often expressed with low abundance in mammalian cells, VcircRNAs with the most abundant raw read counts were selected for wet lab confirmation. To confirm the expression of predicted VcircRNAs, a detection pipeline (Fig. 2A) was enacted initially using RNA isolated from A549-hACE2 cells infected with the WA.1/2020 strain of SARS-CoV-2. From the predicted breakpoints for each putative VcircRNA junction site, primers were designed to amplify upstream of the junction site during Rolling Circle Amplification (RCA) for cDNA synthesis[7], and sequences spanning the unique junction site during PCR amplification of cDNA products. If the target VcircRNAs are present and RCA occurs, amplification products from PCR should produce bands representing single and multiple copies of the VcircRNA on electrophoresis gels. This confirmation protocol was used for the most abundant VcircRNA predicted by SICILIAN, a large predicted VcircRNA with breakpoints in the 7b ORF (5’) and the nucleocapsid encoding (N) ORF (3’) with 560 log-cpm at 72 hpi, which we have named “circ7b8N” (Fig. 2B).

**Figure 2:**
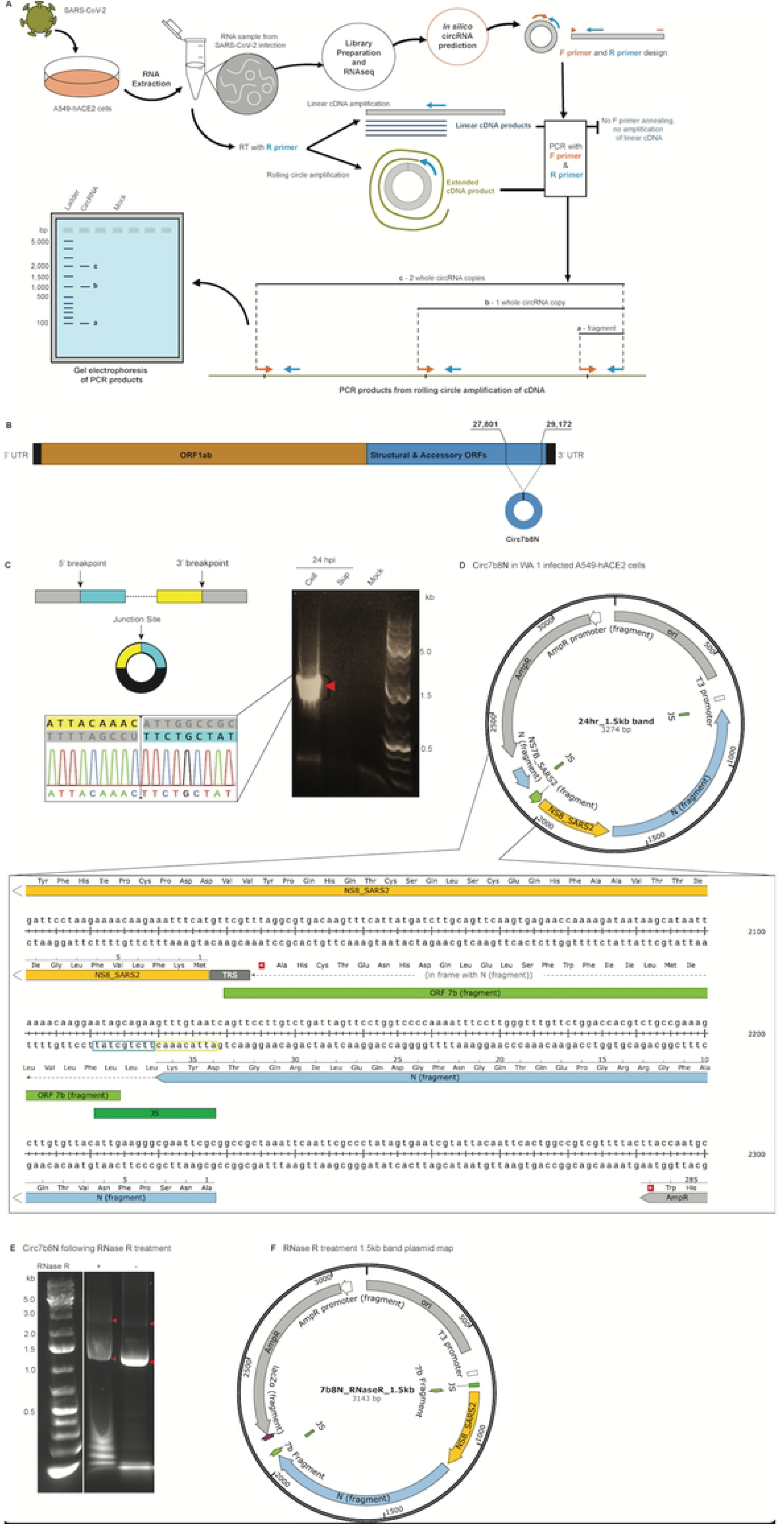
Experimental confirmation of circ7b8N. (A) Experimental workflow for distinguishing circRNA from linear RNA. RNA was isolated from SARS-CoV-2-infected A549-hACE2 cells at 24-, 48-, and 72-hour timepoints post infection, in addition to RNA from uninfected (mock) cells. That RNA was used for VcircRNA predictions via RNAseq and analyses by computational algorithms, as well as for wet lab confirmation of predicted VcircRNAs. If VcircRNAs are present in RNA isolates, they will be identified using rolling circle amplification for cDNA synthesis, followed by traditional PCR amplification using targeted forward (F) and reverse (R) primers which span and flank the unique VcircRNA junction site, respectively. The presence of multiple copies of the VcircRNA sequence along the cDNA can be visually identified by gel electrophoresis. (B) Diagram of the breakpoint locations (top numbers) for circ7b8N, expressed from a combination of structural ORFs and accessory protein ORFs (blue). (C) Agarose gel confirming circ7b8N in infected A549-hACE2 cells (“Cell”) 24-hours post infection (hpi), but not in supernatant (“Sup”) or mock (“Mock”) infected cells, by rolling circle amplification and PCR. Bands indicated with red arrows were excised and DNA extracted for sequencing confirmation (DNA electropherogram). Junction site breakpoints (dashed lines) are shown between breakpoint junction sequence (yellow) and 5’ breakpoint junction sequence (aqua), with upstream and downstream linear sequences shown in grey. (D) Whole plasmid sequencing map of the excised 1.5kb band from the electrophoresis gel in panel B, which was cloned into a TOPO vector prior to sequencing. Two copies of the junction site (neon green) are shown flanking the entire circRNA sequence, with two copies of 7b (olive green), ORF8 (yellow), and N fragment (light blue) inserted into the TOPO vector sequence (grey). Snapshot of sequence extracted from SnapGene. (E) Agarose gel confirming circ7b8N in WA.1 infected cells resists degradation by RNase R treatment. The 1.5kb band was extracted and cloned into a TOPO vector prior to whole plasmid sequencing (F). The ladder lane of this gel was run on the same gel as the samples, but other samples representing another VcircRNA that is not discussed in this paper were removed. White band between the ladder and sample lanes was left to indicate the alteration for transparency. Similar to the plasmid map in D, two copies of the junction site (neon green) and two copies of the 7b fragment (olive green) flank the ORF8 (yellow) and N fragment (light blue) sequences.

Circ7b8N spans 1,372 nucleotides from positions 27,801 to 29,172 in the SARS-CoV-2 genome, encompassing the 3’-most 66% of the 7b ORF sequence, the entire 8 ORF, and the 5’-most 71% of N ORF (Fig. 2B). circ7b8N was only predicted by SICILIAN, not by other algorithms, although other algorithms identified similarly sized VcircRNAs with breakpoints in the 7b and N regions. Computational prediction suggested that circ7b8N was expressed in positive-sense and negative-sense orientations, which we confirmed by using sense-specific designed PCR primers to amplify circ7b8N from SARS-CoV-2 infected A549-hACE2 human lung epithelial cells (Fig. 2A). Circ7b8N was also shown to be resistant to degradation by RNase R (Fig. 2E-F), a unique attribute of circRNAs, yet the resistance varies for each circRNA. Circ7b8N appears to be less resistant to RNase R degradation as other circRNAs, such as those generated by HCV[7] which are highly resistant to RNase R degradation and therefore selected by RNase R treatment. Topoisomerase (TOPO) clones generated using the DNA extracted from gel electrophoresis bands of the appropriate sizes (multiples of ∼1,372 nucleotides) confirm the circular permutation of circ7b8N by the inclusion of at least two copies of the junction site in the circ7b8N gel DNA insert (Fig. 2D, F). While we were unable to see more than two copies of the junction site in the confirmatory sequencing results due to the large size of circ7b8N, the presence of two junction sites with 7b ORF sequences beyond the immediate junction site shown connected to N ORF without interruption suggested adequate rolling circle amplification, and therefore, that circ7b8N is indeed a bona fide circRNA.

Our confirmed VcircRNA is circular, in the sense that it lacks 5’ and 3’ terminal ends, resulting in a continuous loop of RNA. The continuous loop structure confers stability over linear RNA, in that there are no termini or loose ends that could initiate degradation by riboexonucleases. However, predicted secondary structures suggest configurations that are far more complex than a traditional circular “O” loop. By using RNAfold WebServer to assess the likelihood of secondary structures in circ7b8N, we observed numerous, complex secondary structures in circ7b8N (Fig. S3). Secondary structure predictions have yet to be correlated with any circRNA predictive success, in that there are no secondary structure signatures of confirmed circRNA. Rather, structural predictions are suggestive of different folding dynamics, stability or function, and are dependent on the nucleotide sequence of each unique circRNA.

### Detection of circ7b8N in clinical samples

#### Nasopharyngeal swabs

RNA samples were obtained from nasopharyngeal (NP) swabs collected during acute SARS-CoV-2 infection by the Peginterferon Lambda 1a clinical trial study at Stanford University School of Medicine[23,24]. Initial assessment of each RNA sample by reverse transcriptase-polymerase chain reaction (RT-PCR) and sanger sequencing was performed by the Stanford Health Care Clinical Virology Laboratory and has been described previously[23,24]. A collection of 12 confirmed positive RNA samples were obtained and screened for circ7b8N (Fig. 3A). Three samples contained circ7b8N and were confirmed by whole plasmid sequencing (Fig. 3A, Fig. S4). Each of the NP swabs were obtained from acute infections with different variants. Circ7b8N was confirmed from variants XBB.1.16.6, an emergent variant with limited circulation in late summer 2023, FU.1, a variant which emerged in late summer/early fall 2023 with some overlap with XBB1.16.1, and XBB.1.9.1, a variant from Winter 2023 that had an extensive global spread. The nine other samples that tested negative for circ7b8N were from Pango lineages FE.1.2 (Spring 2023), XBB.1.16 (Spring-Summer 2023), EG.1 (Summer 2023), XBB.2.3.11 (Summer-Fall 2023), EG.5.1 (Fall 2023), and XBB.1.9.1 (Winter 2023).

**Figure 3:**
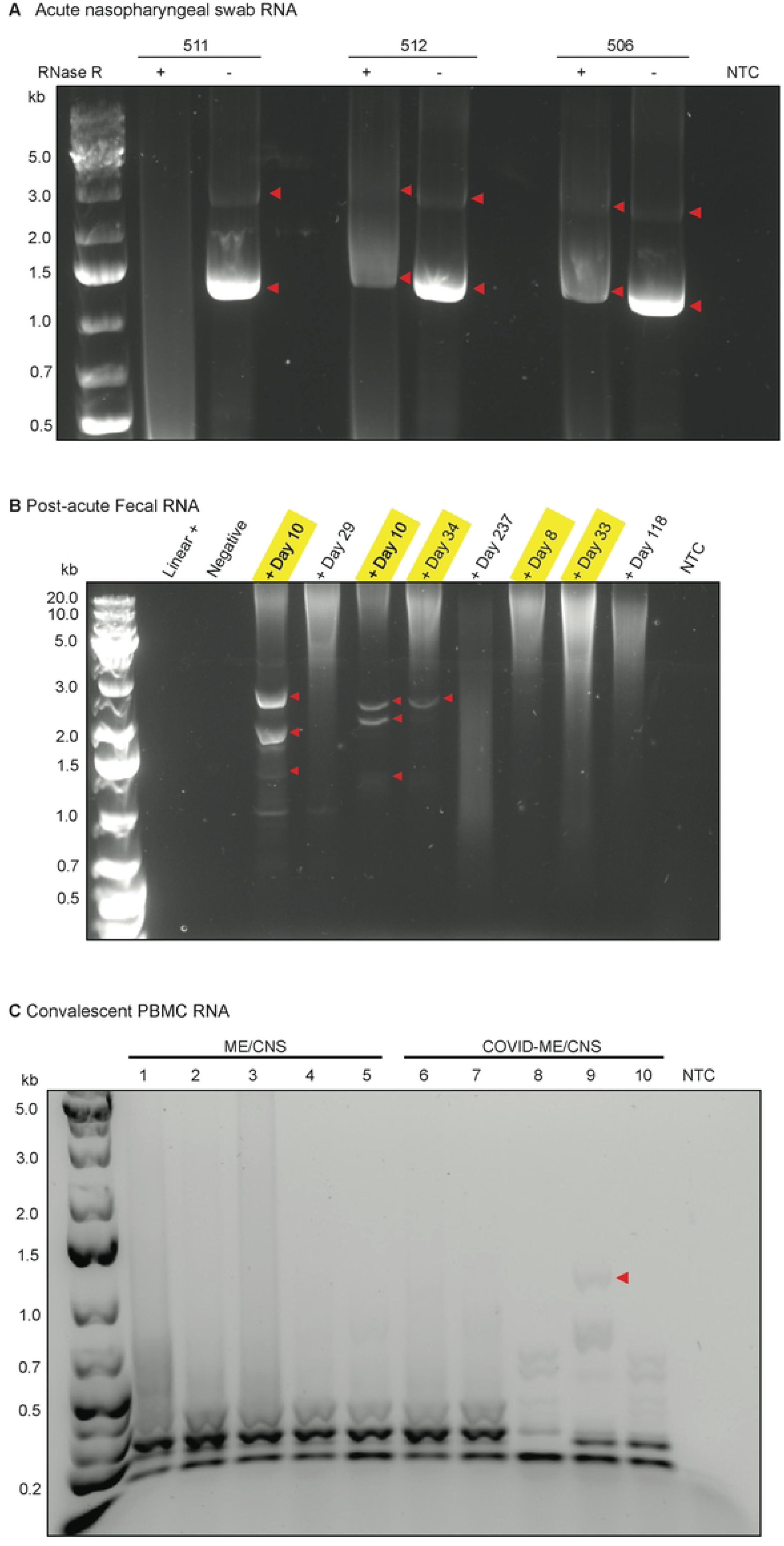
Detection of circ7b8N in clinical samples. RNA eluted from nasopharyngeal (NP) swabs, fecal samples, and peripheral blood mononuclear cells (PBMCs) were screened for the presence of circ7b8N by rolling circle amplification and PCR. (A) Treatment with RNase R was used on NP swab RNA to enrich for potential VcircRNAs. (B) Fecal samples were collected post-acute infection to screen for persistence of circ7b8N. Each lane indicates a sample from a different donor collected at the timepoint indicated in the lane labels. Those with detectable N1 sequences, based on CDC primers, are highlighted in yellow. (C) PBMCs were collected from individuals with myalgic encephalomyelitis/chronic fatigue syndrome (ME/CFS) or ME/CFS associated with COVID-19 infection (COVID-ME-CNS). NTC = no template control.

#### Fecal Samples

Fecal samples collected as a part of the Peginterferon Lambda 1a study were used to detect SARS-CoV-2 beyond NP swabs and monitor viral shedding during post-acute periods[25]. Initial testing of RNA from fecal samples was performed by PCR using standard SARS-CoV-2 diagnostic primer sets from the Center for Disease Control (CDC), as previously described[25]. Of the two primer sets (“N1” and “N2”) designed to amplify 100 nucleotide stretches of the N ORF, the N1 target region is within circ7b8N. Fecal samples which tested positive using the N1 CDC primer set in initial testing also contained circ7b8N by PCR of cDNA produced by RCA (Fig. 3B). Controls for this testing included samples from an uninfected donor and a sample from a fecal sample that had SARS-CoV-2 RNA spiked in, meaning the linear genome was present but was not replicative. Both the uninfected and spiked controls were negative for circ7b8N.

#### Peripheral Blood Mononuclear Cells from Long COVID cases

To investigate whether circ7b8N is associated with progressive COVID symptoms after the clearance of acute infection, or “Long COVID”, peripheral blood mononuclear cells (PBMCs) from five individuals with myalgic encephalomyelitis/chronic fatigue syndrome (ME/CFS) associated with COVID infection, and five individuals with non-COVID-associated ME/CFS were obtained from the Stanford University Post-Acute COVID-19 Syndrome Clinic and screened for circ7b8N using RCA as described earlier. Albeit with a faint band at ∼1,400 nucleotides, this implies circ7b8N may present in sample 9 (Fig. 3C), one of the COVID-associated ME/CFS cases. However, the amount of DNA that could be extracted from this gel was not sufficient for confirmation by sequencing.

### Cellular localization of circ7b8N

The limits of VcircRNA functionality are directly related to where the VcircRNAs exist in the cellular environment. To investigate where circ7b8N is localized in the cellular environment, we performed Fluorescent *In Situ* Hybridization (FISH) in A549-hACE2 cells (1) transfected with synthesized circ7b8N, or (2) infected with the WA.1 strain of SARS-CoV-2 (Fig. 4). A synthesized circRNA encoding eGFP with the coxsackievirus B IRES sequence served as a circRNA negative control. Our probe designed against the junction site of circ7b8N was detectable in cells treated with synthesized circ7b8N and infected with SARS-CoV-2/WA.1/2020, but not in mock/untransfected cells, cells transfected with eGFP circRNA, nor cells transfected with the non-infectious SARS-CoV-2 replicon RNA. In all conditions where circ7b8N was detectable, circ7b8N was detected in punctate structures that are restricted to the cytoplasm, and was not observed in the nucleus, suggesting that it has functional roles in the cytoplasm

**Figure 4:**
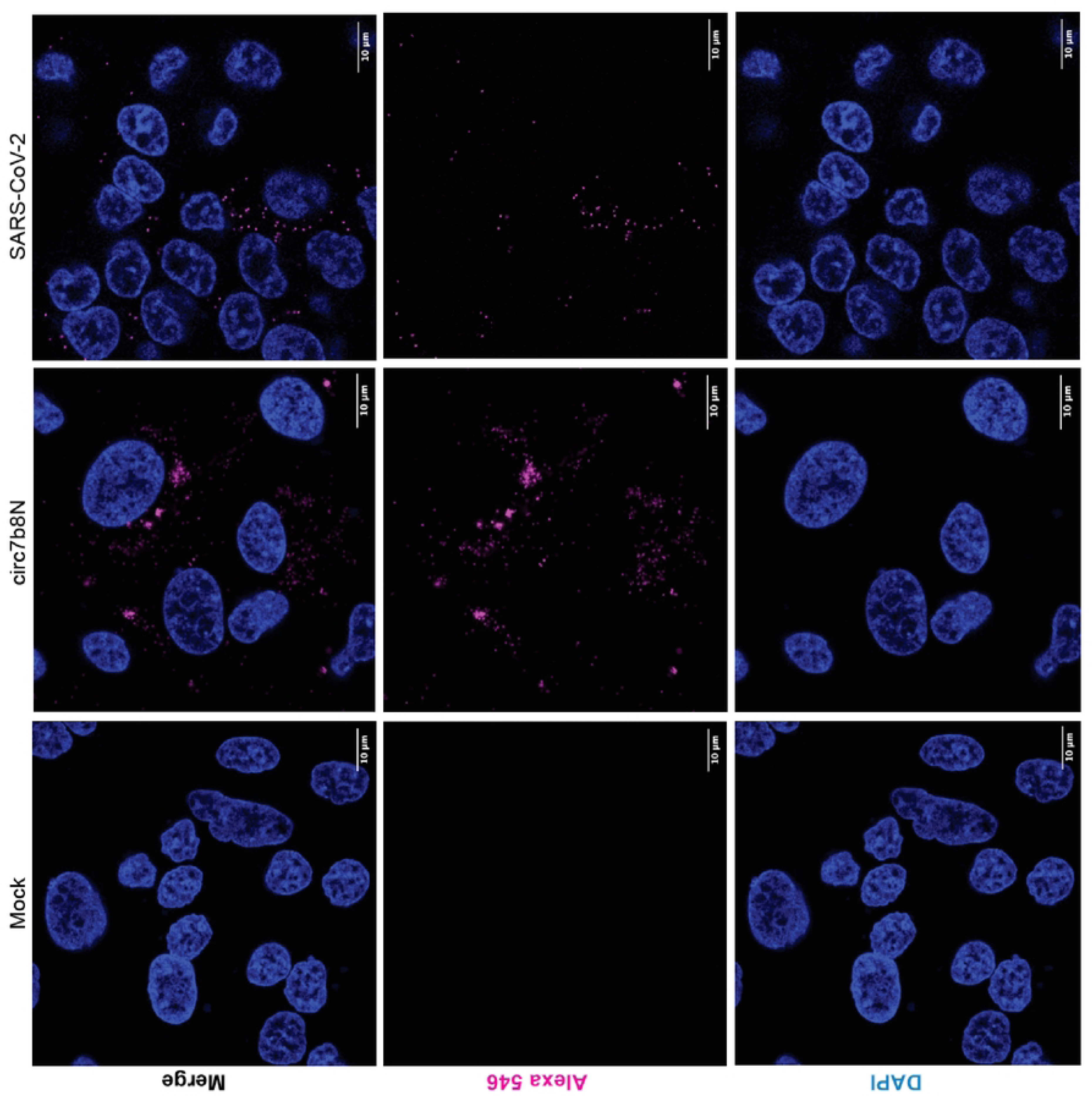
Cellular localization of circ7b8N. Fluorescent In Situ Hybridization (FISH) was performed to identify cellular localization of circ7b8N across infection or transfection type and imaged using confocal microscopy. Blue = nuclei (DAPI), magenta = junction site-specific circ7b8N probes (Alexa 546). Scale = 10 μM.

### Viral kinetics

To determine whether overexpression of circ7b8N affects viral replication, RNA yields and viral titers were quantified by plaque forming assay (PFA) and quantitative PCR (qPCR) (Fig. S5). SARS-CoV-2 infected cells transfected with circ7b8N produced titers equivalent to infected cells without additional circ7b8N (Fig. S5A). Similarly, no observable differences in quantities of the envelope (E) sequence were noted between A549-hACE2 cells with or without added synthesized circ7b8N prior to SARS-CoV-2 infection (Fig. S6B). These results suggest that overexpression of circ7b8N does not impact viral replication kinetics or overall viral yield during SARS-CoV-2 infection, at least not in infected cell cultures. It is possible that VcircRNA 7b8N is already saturated in infected cells to perform its function on viral gene expression, or that its function involves the modulation of host cell gene expression.

#### *Lineage analysis* – expression across variants of concern (VoCs)

Multisequence alignments (MSAs) were performed on representative sequences compiled from each variant of concern (VoC) to assess whether circ7b8N is conserved across variants that were indicated to have a high probability of increased transmission, increased disease severity, and increased escape from antibody memory and available treatments (Fig. 5A, Suppl. Table 1). The 3’ breakpoint and junction region of circ7b8N was found to be entirely conserved across variants that emerged between 2019 and 2024 VoCs (Fig. 5A). Aside from a single point mutation in the 5’ junction region of the 2020 *Iota* variant (nucleotide position 27,804, C to T) that is not carried over into later variants, there are two mutations that do carry over consistently – one with the emergence of the first omicron variant in 2021 (nucleotide position 27,807, C to T), and another with the emergence of later omicron variants in 2024 (nucleotide position 27,810, T to C). Analysis beyond the junction site region for circ7b8N was not conducted, as the 7b and 8 ORFs are known for accumulating mutations, and these are likely significant for translation from the linear genome rather than in the formation of VcircRNA.

**Figure 5:**
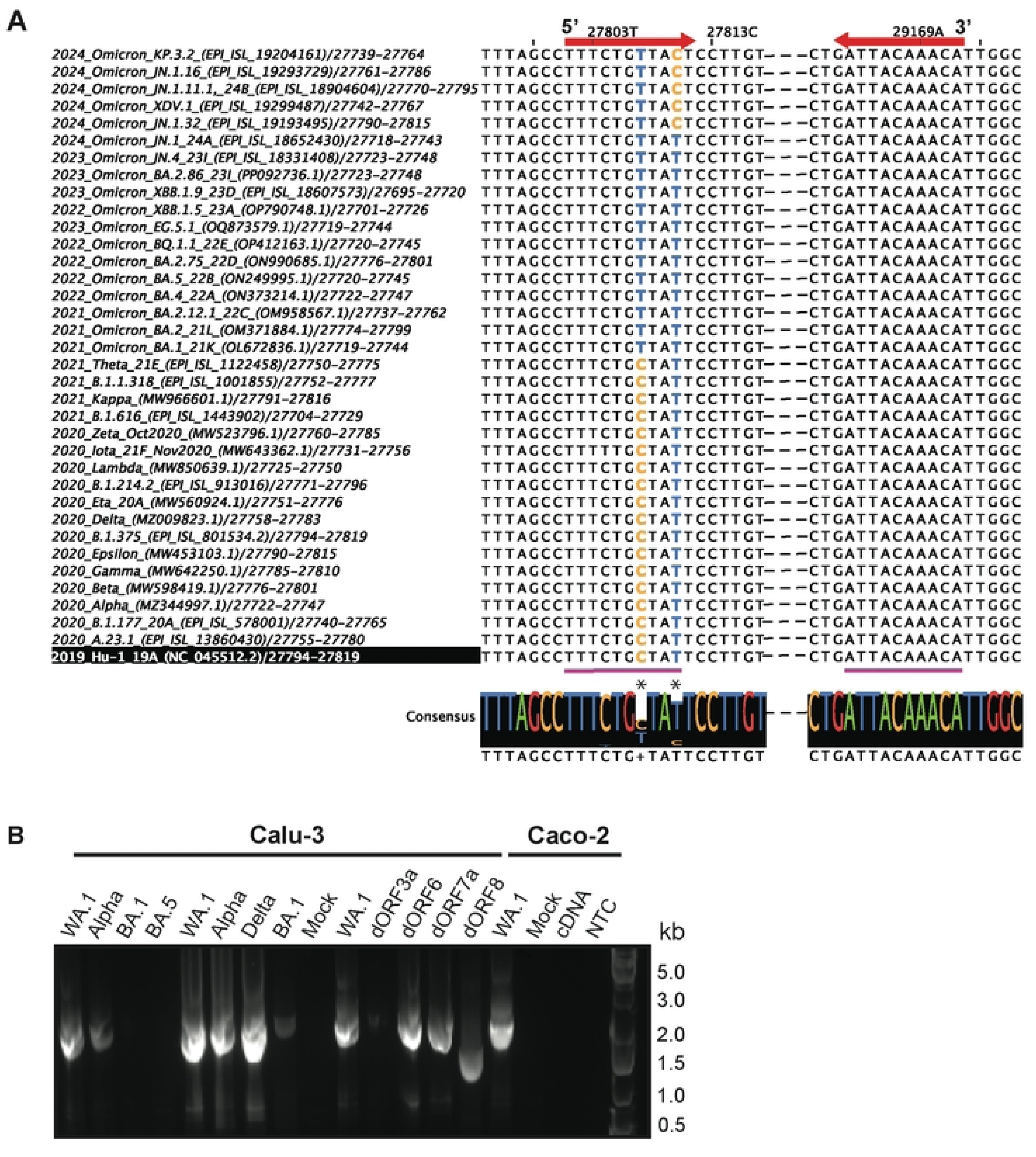
Conservation of circ7b8N across SARS-COV-2 Variants of Concern (VoCs). (A) MSA of VoCs of junction site breakpoints for circ7b8N. Consensus sequence below the alignment composite indicates positions of point mutations (*) within the junction site (pink lines). Red arrows at the top of each alignment indicate the direction of the circRNA sequence from each breakpoint (blunt end of arrows). The 2019 Hu.1 strain (highlighted in black, bottom) was the reference sequence. Library preparations for RNAseq convert Uracil (U) to Thymine (T), as reported here. (B-C) Confirmation of circ7b8N expression in Calu-3 and Caco-2 cell lines using different SARS-CoV-2 variants. Calu-3 or Caco2 cells were infected with indicated SARS-CoV-2 WA1, Alpha, and Omicron variants (BA.1 and BA.5) with 0.1 MOI and 48 h later, total RNA was collected. Circ7b8N was confirmed by sequencing in all variants, including Omicron variants BA.1 and BA.5, which required 2x PCR processing for band visualization. Samples indicated by “dORF” are infected with mutant WA.1 strains with deleted ORF3a, ORF6, ORF7a, or ORF8.

Next, Calu-3 cells were infected with SARS-CoV-2 strains WA.1 (circulating 2020), Alpha (circulating Spring 2020 – Summer 2021), Delta (circulating Summer 2020 – Winter 2021), Omicron BA.1 (circulating Spring – Fall 2021), Omicron BA.5 (circulating Winter 2021 – Spring 2022), and Caco-2 cells were infected with WA.1. Circ7b8N was confirmed by PCR (Fig. 5B) and whole plasmid sequencing in all infections (Suppl. Fig. S6), although infections with Omicron BA.1 and BA.5 required two rounds of PCR for visible banding. Additionally, circ7b8N was found to be present in generated strains of WA.1 to contain single, whole-ORF deletions from ORF 3a (dORF3a), ORF 6 (dORF6), ORF 7a (dORF7a), and ORF 8 (dORF8). Of the WA.1 deletion mutants, all were positive for circ7b8N expression, and dORF8 showed circularization with the sample circ7b8N junction site, yet with a truncated sequence containing only 7b and N fragments from the original circ7b8N predicted sequence. This suggests that VcircRNAs may be expressed with large deletions, and likely mutations, to their internal sequence, if the junction site is present.

### Lineage analysis – expression across CoVs

Multisequence alignment (MSA) and a phylogenetic tree were constructed using a compiled list of CoV complete genomes to assess conservation of circ7b8N across *Coronaviridae* species (Fig. S7) (Suppl. Table 2). Emphasis was placed on junction site breakpoint conservation to confirm the likelihood that circ7b8N could be expressed by other CoVs. Given the large length of circ7b8N, variations within the VcircRNA sequence beyond the junction site were not analyzed, as any variation would likely not be unique from linear transcripts. Variability across CoVs was significant, with little-to-no conservation of circ7b8N junction sites beyond BANAL viruses (Fig. S7A). MSAs were analyzed for sequence motifs of circ7b8N junction sites and for positionality along the genome (i.e. specific nucleotide region corresponding with each junction site break point), as many of the non-human genomes were minimally annotated. This suggests that the emergence of circ7b8N as a distinct VcircRNA likely occurred as SARS, SARS-CoV-2 and BANAL lineages diverged from other bat CoVs and their common ancestors in the beta-CoV genus (Fig. S7B).

### Effects of VcircRNA 7b8N on host gene expression

To assess the effects of circ7b8N on host cell environments, we analyzed differential gene expression in cells transfected with VcircRNA 7b8N on host gene expression (Fig. 6-7). All experimental conditions were performed in triplicate and variation between replicates, as indicated by Spearman correlation values, was low. A total of 108 and 435 differentially expressed genes (p < 0.05) were identified in circ7b8N-treated samples in comparison to mock (Fig. 6) and SARS-CoV-2-infected (Fig. 7) samples, respectively. GO term calls were trimmed for redundancy and minimum fold change of >1 to represent the most enriched genes in circ7b8N-treated cells (Figs. 6B, 7B). Thresholds of <0.001 or <0.05 False Discovery Rate (FDR) scores were applied to DESeq2 results prior to gene ontology (GO) term analyses.

**Figure 6:**
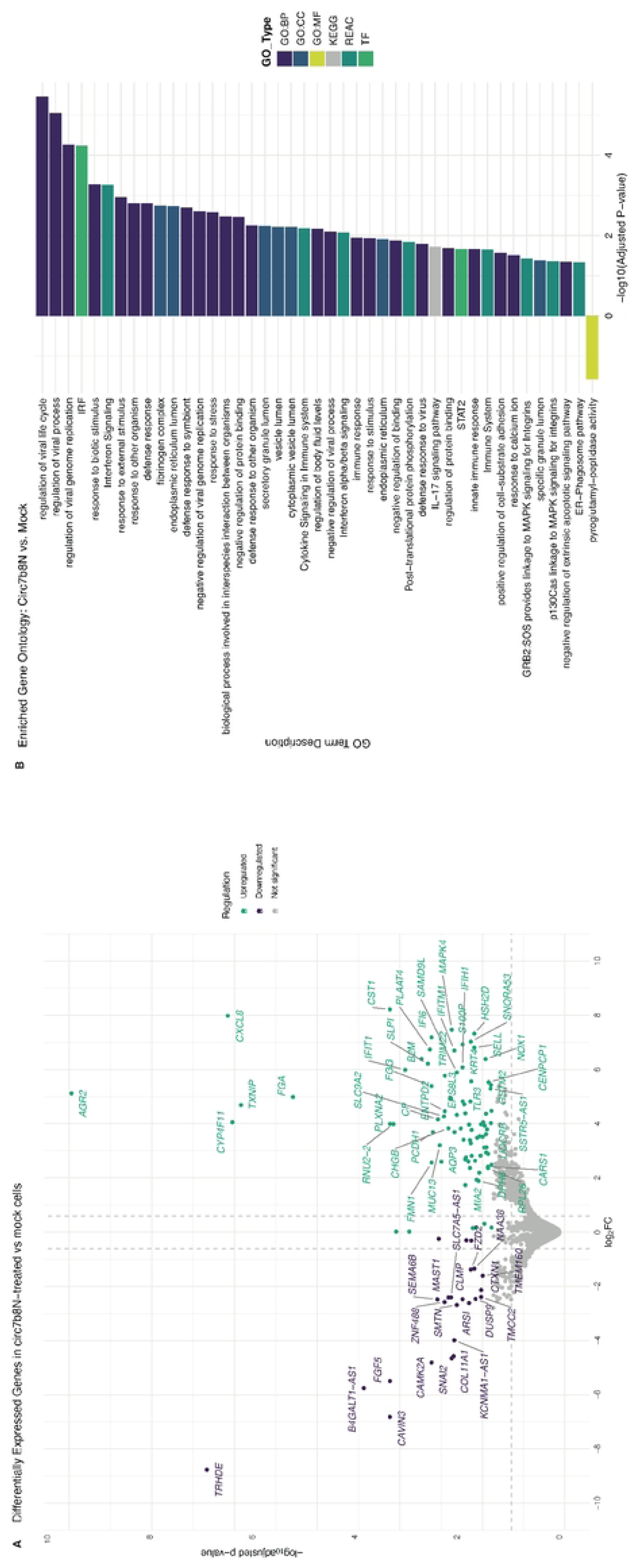
Differentially expressed genes: circ7b8N vs. Mock. (A) Volcano plot of significantly differentially expressed genes in A549-hACE2 cells after treatment with circ7b8N. Top 30 significantly up- and down-regulated genes are labeled and were determined by a p-value cutoff of 0.05. (B) Gene ontology terms of differentially expressed genes in response to circ7b8N treatment. GO terms were generated from a list of differentially expressed genes with an FDR of <0.01 compared to mock/untransfected A549-hACE2 cells. GO:BP=Biological Process; GO:MF=Molecular Function; KEGG=Kyoto Encyclopedia of Genes and Genomes; REAC=Reactome; TF= Transcription Factor.

**Figure 7:**
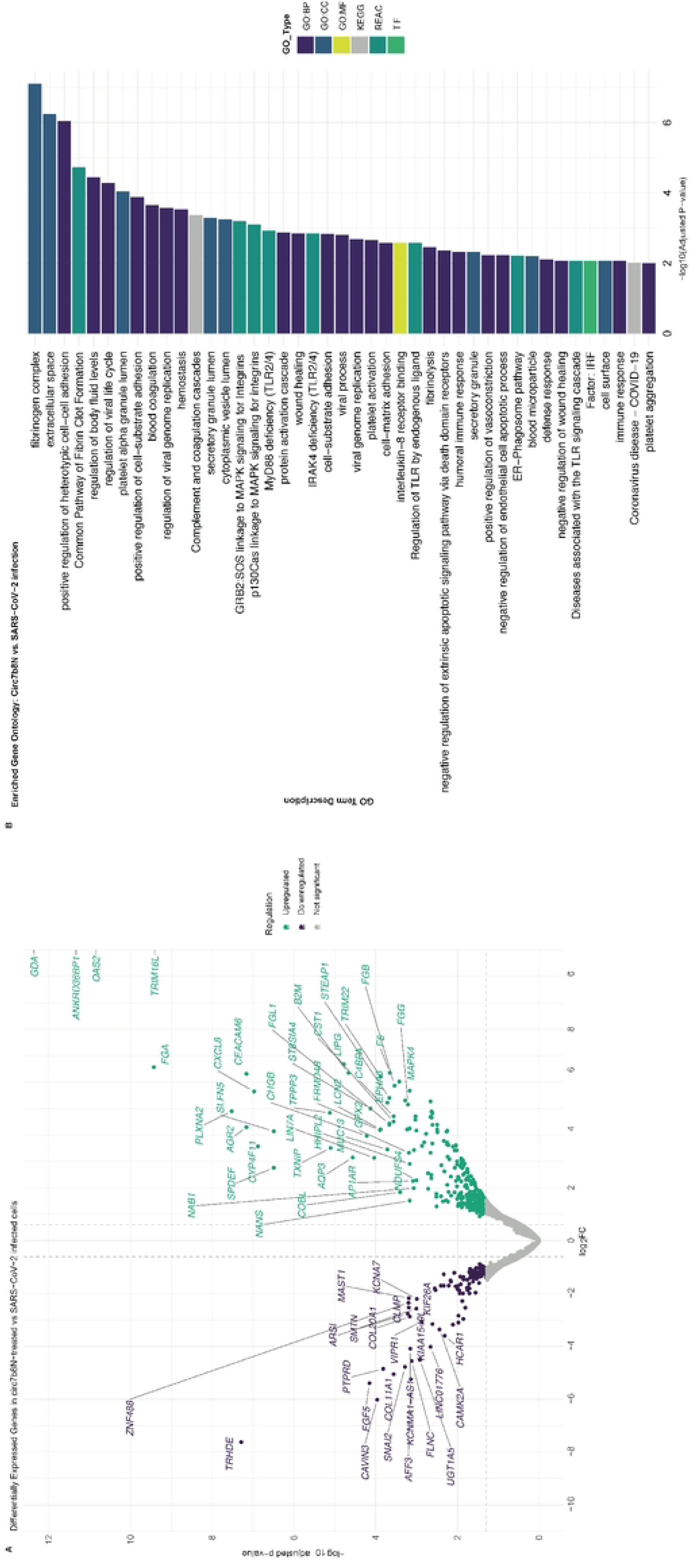
Differentially expressed genes: circ7b8N vs. SARS-CoV-2 infection. (A) Volcano plot of significantly differentially expressed genes in A549-hACE2 cells after treatment with circ7b8N. Top 20 significantly up- and down-regulated genes are labeled and were determined by a p-value cutoff of 0.05. (B) Gene ontology terms for host genes that are differentially expressed in response to circ7b8N compared to SARS-CoV-2 infection. GO terms were generated from a list of differentially expressed genes with an FDR of <0.01 compared to A549-hACE2 cells infected with SARS-CoV-2. GO:BP=Biological Process; GO:CC=Cellular Component; GO:MF=Molecular Function; KEGG=Kyoto Encyclopedia of Genes and Genomes; REAC=Reactome; TF= Transcription Factor.

Compared to non-transfected A549-hACE2 cells, transfection of circ7b8N into A549-hACE2 cells significantly enriched 84 genes (p_adj_ < 0.05), with the top including Anterior Gradient 2 (AGR2), Interleukin-8 (CXCL8), Cytochrome P450 family 4 subfamily F member 11 (CYP4F11), Thioredoxin-interacting protein (TXNIP), Fibrinogen Alpha Chain (FGA), Cystatin SN (CST1), Plexin-A2 (PLXNA2), RNA U2 small nuclear 2 functional noncoding RNA gene (RNU2-2), Interferon-Induced Protein with Tetratricopeptide Repeats 1 (IFIT1), and Tripartite Motif Containing 16 Like pseudogene (TRIM16L, also known as TRIM70) (Fig. 6A, green, Suppl. Table 3). Additionally, 24 genes were significantly suppressed or downregulated (p_adj_ < 0.05) by circ7b8N compared to mock condition samples, including Thyrotropin-Releasing Hormone Degrading Enzyme (TRHDE), Caveolae Associated Protein 3 (CAVIN3), Calcium/Calmodulin-Dependent Protein Kinase II Alpha (CAMK2A), Snail Family Transcriptional Repressor 2 (SNAI2), Collagen Type XI Alpha 1 Chain gene (COL11A1), Zinc Finger Protein 488 (ZNF488), Semaphorin 6B (SEMA6B) and Microtubule Associated Serine/Threonine Kinase 1 gene (MAST1) (Fig. 8A, purple, Suppl. Table 3). Genes such as B4GALT1-AS1 and FGF5 are likely genes important for cell line immortalization, yet their downregulation might represent cell stress from the transfection and warrants further investigation.

**Figure 8:**
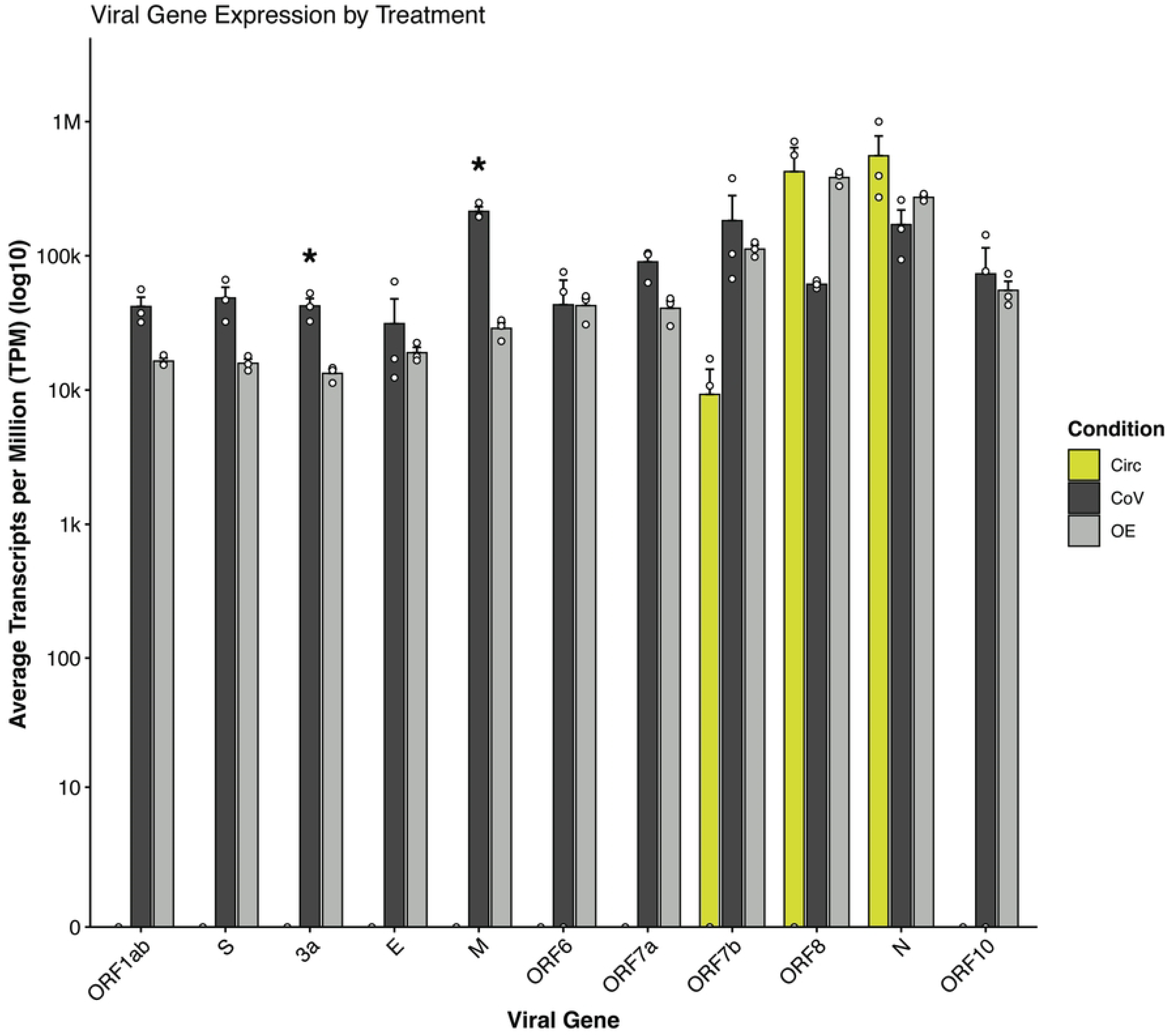
SARS-CoV-2 Differential Gene Expression. Circ = circ7b8N-only transfection, CoV = infection only, OE = circ7b8N+infection. Treatments and sequencing for each were performed in independent triplicate experiments. The presence of ORFs 7b, 8 and N in circ7b8N-treated samples (Circ, yellow bars) compared to infected samples and overexpression samples are due to the transfection of circ7b8N. Error = Mean + SEM, * = p < 0.05 (one-way ANOVA (for ORF7b, ORF8, and N), or t-test).

Gene ontology terms associated with genes enriched by treatment with circ7b8N broadly represent biological processes (GO:BP) and cellular components (GO:CC) related to immune response to infection and regulation of viral life cycle processes compared to mock samples (Fig. 6B). Specifically, chemokine CXCL8, Secretory Leukocyte Protease Inhibitor 1 (SLPI), and core interferon stimulating genes IFITM1, IFIT1, IFI6, and TRIM22 were all significantly enriched (p_adj_ < 0.000003), suggesting circ7b8N stimulates pro-inflammatory antiviral responses (GO term “Regulation of viral life cycle” (-log10 p_adj_ = 5.46) (Fig. 8B, Suppl. Table 4). Genes associated with cellular components were significantly enriched, including components supporting scaffolding within the extracellular matrix (fibronectin complex (-log10 p_adj_ = 2.75)), protein and lipid synthesis (endoplasmic reticulum lumen (-log10 p_adj_ = 2.73)), and vesicle formation and trafficking (secretory granule lumen (-log10 p_adj_ = 2.24) and cytoplasmic vesicle lumen (-log10 p_adj_ = 2.21)), all of which are cellular structures and locations exploited by SARS-CoV-2[26,27]. Interferon Regulatory Factor (IRF) and Signal Transducer and Activator of Transcription 2 (STAT2), known transcription factors, were also enriched, suggesting an enhanced interferon response to circ7b8N.

In contrast to SARS-CoV-2-infected cells (Fig. 7), treatment with circ7b8N alone, without viral infection (Fig.6), resulted in significant (P_adj_ < 0.01) enrichment of 101 genes and suppression of 34 genes. Significantly enriched genes include Guanine Deaminase (*GDA*, P_adj_ = 7.87E-13), Ankyrin Repeat Domain 36B Pseudogene 1 (*ANKRD36BP1*, P_adj_ = 6.06E-12), 2’-5’-oligoadenylate synthetase 2 gene (*OAS2*, P_adj_ = 1.10E-11), Carcinoembryonic Antigen-Related Cell Adhesion Molecule 6 gene (*CEACAM6*, P_adj_ = 6.80E-8), SAM Pointed Domain Containing ETS Transcription Factor gene (*SPDEF*, P_adj_ = 3.19E-7), and Schlagen Family Member 5 gene (*SLFN5*, P_adj_ = 3.19E-7). Genes significantly suppressed by circ7b8N treatment included Protein Tyrosine Phosphatase Receptor Type D (*PTPRD*, P_adj_ = 0.00015) and AF4/FMR2 Family Member 3 gene (*AFF3*, P_adj_ = 0.0007). Genes such as *TRIM16L*, *B2M*, *FGA*, *TXNIP*, *CYP4F11*, and *PLXNA2* were significantly enriched by circ7b8N treatment in comparison to both mock and infected cells. Similarly, *TRHDE*, *FGF5*, *CAVIN3*, and *SNAI2* were also downregulated in both comparative analyses.

Transfection with circ7b8N alone enriched specific pathways related to membrane and cell-cell adhesion and coagulation, suggesting circ7b8N may play a role in autophagy[28], or influence broader tissue- and organ-level symptoms during infection. Differentially expressed genes relating to coagulation, clot formation, hemostasis, and body fluid levels were prominently enriched in cells transfected with circ7b8N compared to SARS-CoV-2 infected cells, further supporting the role of circ7b8N in pathways with larger, systemic impacts. Specifically, Fibrogen α-chain (*FGA*) is repeatedly represented and highly enriched (-log10 p_adj_ = 7.10, Suppl. Table 4) in our dataset, as others have observed with similar significance in SARS-CoV-2-infected tissues[29].

Circ7b8N exposure also enriched genes with finer granularity, such MAPK signaling and IL-8 receptor binding, which indicate a heightened pro-inflammatory response[30]^,[31]^. *OAS2*, an innate immune response protein that detects double stranded RNA and triggers RNA degradation by RNAse L and an antiviral cascade[32], was only enriched in circ7b8N treatment conditions when compared to SARS-CoV-2 infection (Fig. 7). While circ7b8N stimulated antiviral responses when compared to genes expressed in mock conditions, *OAS2* and other viral RNA-binding proteins were not enriched by circ7b8N unless analyzed against SARS-CoV-2 infected cells. Signatures of secretory granules and vesicle formation are also enriched, indicating circ7b8N may be involved in regulatory functions for replication organelle formation and support, viral assembly and trafficking, and egress.

### Influence on SARS-CoV-2 gene expression

In addition to modulating host gene expression, we wondered whether circ7b8N acted on viral genomic and sgRNA to modulate SARS-CoV-2 viral gene expression during infection. Here we compare SARS-CoV-2 gene expression across circ7b8N-treated cells (“Circ”), SARS-CoV-2-infected cells (“CoV”), and circ7b8N-treated cells infected with SARS-COV-2 (OE, or “overexpression”) (Fig. 8). Notably, a significant decrease in ORF3a and ORF M expression was observed in the OE condition compared to infected cells without the addition of circ7b8N (CoV). Decreases in ORF1ab, ORF S and ORF 7a were also observed, whereas ORF 6 and ORF 10 remained static across infected and overexpression conditions. Based on our findings, it is unlikely that these differences in viral gene expression are due to circ7b8N acting directly on the viral genome directly, although further experimentation is required to confirm mechanisms of action.

## Discussion

This work confirms that circ7b8N, a distinct VcircRNA molecule is expressed from the SARS-CoV-2 viral genome. While we could not deplete the VcircRNA, overexpression of circ7b8N did not display any effects on viral replication, suggesting that circ7b8N influences SARS-CoV-2 infection dynamics through impacts on the host cellular environment. This is evident with our ability to detect circ7b8N consistently across cell culture and clinical samples during acute infection and with the quantification of differential host gene expression in response to circ7b8N exposures. Localization of circ7b8N outside of the nucleus during infection indicates circ7b8N is generated and utilized in the cytoplasmic environment, likely in the double membrane vesicles (DMVs) which support viral replication[3,4,33–35], rather than in the nucleus, where host endogenously-expressed circRNAs are synthesized. Our data showing that circ7b8N localizes to the cytoplasm does not exclude circ7b8N trafficking and transfer between cells, which has been previously observed with exogenous circRNAs[36] and is commonly exploited for biomedical biomarker applications[37–39]. Our transfection conditions were designed to simulate uninfected bystander cells receiving circ7b8N and thus illustrate stimulation of host antiviral responses agnostic of DMV formation. Future experiments are needed to distinguish between circ7b8N molecules which are trafficked to uninfected bystander cells and those which are expressed during replication in DMVs to better understand whether the method of introduction of circ7b8N shapes distinct functionality.

Our interrogation of differentially expressed host genes in response to circ7b8N provided evidence to suggest that VcircRNAs can stimulate an immune response via the enrichment of many interferon stimulating genes. Specifically, signatures of apoptosis and autophagy were significantly enriched, and thus further experimentation is warranted to identify the specific roles circ7b8N may play in these important defense responses. For example, we observed significant enrichment of CXCL8 by circ7b8N treatment, suggesting circ7b8N alone can stimulate chemokine responses. Others have found the early expression of CXCL8 in response to EV-D68, rhinoviruses, and influenza virus enhances viral replication in respiratory cells[40]. TXNIP, a protein which activates inflammasome complex formation in the endoplasmic reticulum and promotes inflammation and apoptosis[41], was enriched by circ7b8N treatment. Overexpression of TXNIP promotes oxidative stress responses by increasing production of reactive oxygen species[41,42]. In other studies, TXNIP was shown to be downregulated in response to Omicron BA.1 infection, which not only differs from our findings, but does not correlate with our findings that circ7b8N is widely conserved across SARS-CoV-2 variants. This may be due to differences in protocol, as Rong et al. utilized hACE2 transgenic mice to analyze systemic infection and TXNIP downregulation was only reported in goblet cells, smooth muscle cells, fibroblasts, and capillary endothelial cells[42], whereas our differential gene expression experiments were limited to hACE2-A549 cells in culture. *CST1*, which regulates peptide bond cleavage and inhibits cysteine proteases in inflammatory airway diseases and acts as an early inhibitor to many viral infections[43], was found to be enriched by circ7b8N. *TRHDE*, a gene and antisense long noncoding RNA (lncRNA) that sponges miRNA in the cytoplasm and is associated with hepatitis B virus (HBV) prognoses[44] and HIV-affiliated hepatic fibrosis[45], was found to be significantly downregulated in comparison to the mock condition, representing a suppression of pyroglutamyl-peptidase activity.

While we discovered the expression of some genes to be similarly influenced by circ7b8N regardless of comparison group, the comparisons to mock and infected cells describe widely different response landscapes to circ7b8N, which may suggest circ7b8N functions can vary in infected cells or bystander cells. For example, IFIT1, known for its roles sensing viral RNA and inducing antiviral responses, was notably enriched only when compared to mock cells, rather than when compared to SARS-CoV-2 infected cells, despite circ7b8N lacking the 5’ or unmethylated cap required for IFIT1 binding. The SARS-CoV-2 genome has been shown to be highly structured[46,47], but the fragmentation of the viral genome as circRNAs are generated may alter the folding or binding restraints and create unique structures that are not present in the complete, linear genome. Such secondary structures introduce the possibility for double-stranded RNA detection, and activation of downstream antiviral responses[48]. Endogenous circRNAs have been predicted to regulate immune responses, broadly [9–11,48] and specifically in response to SARS-CoV-2[49], therefore specific VcircRNAs, such as circ7b8N, may represent a complementary adaptation which also shape immune responses.

Other significant gene enrichment suggests that circ7b8N may influence membrane remodeling in organelles that are exploited by SARS-CoV-2 for the formation of replication organelles and for packaging and egress during later stages of the viral lifecycle. Remodeling of organelles to support viral replication and egress is a well-documented process[34,50–54], but there have been no mentions of circRNAs nor VcircRNAs involved, as they are often overlooked as potential contributors. Alternatively, these signatures may be indicative of heightened cell signaling in response to viral RNA, triggered by just a fragment (circ7b8N) and not the complete viral genome. As RNA can be trafficked between cells, further investigation should explore whether circ7b8N and other VcircRNAs can pre-empt viral infection, possibly by stimulating membrane remodeling or other mechanisms in uninfected, bystander cells.

Additionally, we detected changes in abundance of specific SARS-CoV-2 viral genes in response to circ7b8N overexpression. Two genes, ORF 3a and M, were significantly downregulated in response to additional circ7b8N. Aside from providing the physical membrane structure for new virions, the M protein orchestrates viral assembly by interacting with N proteins and viral RNA[55]. Cryo-electron microscopy analyses of M protein dimers show structural similarity to the viral ion channel formed by 3a[55], which has been shown to mediate apoptosis[56]. If the lower abundance of either of these genes had resulted in a reduced viral replication, we could theorize disruption of viral assembly and viral ion channel functioning by circ7b8N. Rather, the lack of clear pro- or anti-viral effects on SARS-CoV-2 replication may propose a compensatory effect instead, as treatment with circ7b8N enriched host genes that may influence viral replication, assembly, and egress, as well as apoptosis. It is possible that the enrichment of proteins associated with vesicle and membrane lumen organization by circ7b8N may support other known functions by M and 3a, like the formation of dense bodies formed by 3a and its recruitment of M and S proteins to enhance viral infectivity[57]. The 3a dense bodies, reported by Hartmann et al., are also found via homologs in bat and pangolin coronaviruses[57]. Viral egress and release is also mediated by 3a through the promotion of the lysosomal exocytosis pathway[58,59]. It appears there are many opportunities for circ7b8N to support the diverse roles of 3a, thus further experimentation should be dedicated to teasing apart this relationship.

To assess the utility of VcircRNAs towards SARS-CoV-2 viral fitness and evolutionary origins, we screened CoVs across subcategories for conservation of circ7b8N. By analyzing CoV genomes beyond SARS-CoV-2, we aimed to determine whether VcircRNAs follow expression patterns across CoV lineages. That is, could we observe distinct VcircRNAs retroactively to draw conclusions about the origins of SARS-CoV-2 from other CoVs, and if so, were those CoVs limited to certain species, such as bats, or uniquely severe in humans (MERS and SARS)? We also used this approach to assess whether trends of severity across SARS-CoV-2 VoCs were associated with circ7b8N expression. Circ7b8N was found to be conserved with high fidelity across variants. ORFs 7b and 8 have been shown to be particularly prone to mutations in some variants. ORF8 is within a hypervariable region that is known for nucleotide deletions, substitutions, and recombination. The junction site of circ7b8N was found to be highly conserved across VoCs and in specific *Betacoronaviruses* of close lineage (RaTG13 and BANAL strains, as well as SARS). Despite point mutations in the junction site which persist in Omicron and later variants, circ7b8N was consistently identified in multiple variants. The consistent expression of circ7b8N across variants despite the emergence of mutations within the junction site region suggests circ7b8N may provide vital utility beyond variant fitness.

Despite ample evidence to support zoonotic emergence, the origin of SARS-CoV-2 is controversial and has been politically weaponized. There are decades of wildlife surveys, viral challenges, and research that points to bats as the natural reservoir for a broad range of viral families[60–66]. Conservation of circ7b8N in BANAL and RaTG13 strains is further evidence suggesting SARS-CoV-2 emerged naturally from zoonotic origins. Additionally, there is growing evidence to support recombination as a driving force for the emergence of SARS-CoV-2[67,68], yet the conservation of circ7b8N suggests that it is not likely to be a region of recombinatory significance that aided emergence. Our finding that the junction site sequences for circ7b8N are conserved within the lineage that includes SARS proposes an important emergence of circ7b8N as SARS and later BANAL, RaTG13, and SARS-CoV-2 sequences diverge from their ancestral lineage. Our work provides evidence that circ7b8N is likely conserved across other closely phylogenetically-related CoV lineages, suggesting that circ7b8N may have played a supportive role in the emergence of SARS-CoV-2 from related CoVs, and that the conservation of its sequence and expression was potentially advantageous as SARS-CoV-2 evolved from ancestral CoVs.

While the stark contrast in read frequency between predicted circ7b8N and other predicted VcircRNA expressed by SARS-CoV-2 may suggest functional importance, this may also be representative of overall viral abundance in a concentrated cell culture infection, or progressive accumulation of specific VcircRNAs. Other viruses have reported lower predicted VcircRNA frequency. Specifically, HCV generates only ∼70 abundantly VcircRNAs[7]. This difference could be due to the different abundances of viral RNAs in infected cells. Our ability to confirm circ7b8N expression during SARS-CoV-2 infection and the observation of differentially expressed host genes in response to treatment with circ7b8N warrant further investigation to characterize the importance and functionality of circ7b8N during SARS-CoV-2 infection. This work was limited in our ability to investigate the role of circ7b8N through various depletion strategies, as our *many* attempts at depleting circ7b8N by siRNAs and CRISPR Cas13d were unsuccessful. We believe our junction site targets were restricted by secondary structures, and thus our functional analyses presented here are limited to overexpression.

The high-fidelity conservation, along with our findings suggesting modulation of the host immune response, argue that circ7b8N is beneficial to SARS-CoV-2 more broadly, regardless of variant fitness and disease severity. Yet, without a clear mechanism of action, we are wary of overstating its importance. Our findings open the door for deeper interrogation into the biogenesis and functionality of circ7b8N, not only during SARS-CoV-2 infections, but in the cellular environment. Given the diversity of species susceptible to SARS-CoV-2 infection, it may be that differences in cell type and species may elicit different functions for circ7b8N.

## Acknowledgements

This study was supported by NIH grant 1U19AI171421 (Glenn, PI., Sarnow, PI of sub-project #2), and a Stanford University DARE Fellowship awarded to G-S (2023 – 2025). We are grateful to the ongoing support by Dr. Judith Frydman (Stanford University).

## Author Contributions

**Conceptualization:** Elysse Grossi-Soyster, Peter Sarnow

**Funding Acquisition:** Elysse Grossi-Soyster, Peter Sarnow

**Formal analysis:** Elysse Grossi-Soyster, Rebekah Gullberg, Arjun Rustagi, Jae Seung Lee, Catherine Blish, Sara Cherry, Julia Salzman, Peter Sarnow

**Investigation:** Elysse Grossi-Soyster, Rebekah Gullberg, Arjun Rustagi, Jae Seung Lee

**Project administration:** Elysse Grossi-Soyster, Catherine Blish, Sara Cherry, Julia Salzman, Peter Sarnow

**Supervision:** Catherine Blish, Sara Cherry, Julia Salzman, Peter Sarnow

**Writing – original draft:** Elysse Grossi-Soyster

**Writing – review and editing:** Elysse Grossi-Soyster, Rebekah Gullberg, Arjun Rustagi, Jae Seung Lee, Catherine Blish, Sara Cherry, Julia Salzman, Peter Sarnow

## Materials and Methods

### Sample preparation for initial circRNA prediction

Human A549 lung epithelial cells expressing human angiotensin-converting enzyme 2 (hACE2), grown in complete DMEM media with 50 mg/ml Geneticin selective antibiotic (G418 sulfate, #10131035, Thermo Fisher Scientific, Waltham, MA, USA) were infected with the WA.1 strain of SARS-CoV-2. All infections were performed under BSL-3 laboratory conditions. Total RNA was extracted from cells and supernatant from infected cells at 24-, 48-, and 72-hours post infection (HPI). RNA was also extracted from uninfected A549-hACE2 cells as a mock or uninfected control. RNA extractions were performed in the BSL-3 facility prior to being transferred to the BSL-2 facility for downstream analyses. One microgram (mg) of RNA from each condition with an RNA sample index score (RIN) of 10 was dissolved in nuclease-free water and sent to Novogene (Novogene Corporation Inc., Sacramento, CA, USA) for library preparation and RNA sequencing. Libraries were prepared using the NovaSeq PE150 library preparation kits and sequenced on a NovaSeq6000.

### Acquisition of additional sequencing data for computational comparison

Wylver et al. performed RNAseq on Calu3 cells infected with SARS-CoV-2 in May 2020. Total RNAseq FASTA files were acquired through the NCBI Gene Expression Omnibus (GEO) (Accession # GSE148729) using the faster-q dump function of the SRA toolkit (NCBI, https://github.com/ncbi/sra-tools) on the Stanford high performance computing (HPC) cluster, Sherlock. Sequences were trimmed using Trim Galore! (v. 0.5.0, https://www.bioinformatics.babrahame.ac.uk/projects/trim_galore/) to removed Illumina adapter sequences, and assessed for read quality using FastQC (https://www.bioinformatics.babraham.ac.uk/projects/fastqc/). Following trimming and quality assessment, reads were indexed for alignment with STAR and Bowtie2 and aligned to the SARS-CoV-2 reference genome (NCBI Accession #NC_045512.2). The SARS-CoV-2 reference genome was acquired from NCBI (<u>RefSeq assembly GCF_009858895.2</u>).

### Computational prediction of circRNAs expressed by SARS-CoV-2

Analysis of RNAseq data to identify potential VcircRNAs was initially performed using the SICILIAN algorithm (https://genomebiology.biomedcentral.com/articles/10.1186/s13059-021-02434-8), which compiles aligned RNAseq reads to statistically identify splice junction sites. Analyses were performed with alignments to the SARS-CoV-2 reference genome to identify VcircRNAs. The resulting databases of predicted splice sites were sorted by number of raw sequencing reads as a proxy for abundance prior to quantification. Other algorithms specifically designed to detect circRNA junction sites were used to validate the SICILIAN results, including CIRI2, circRNA_finder, and find_circ algorithms. Cross-algorithm results were compared to results published by Cai et al., which used the same source RNAseq files from Wyler et al. Additional analyses were performed using SPLASH, an unbiased algorithm that identifies sequence anomalies and splice sites prior to alignment to a reference genome.

Junction site break points of predicted VcircRNAs were analyzed for possible breakpoint motifs or signatures. Calls of nucleotide strings representing VcircRNAs predicted by each algorithm were organized in Excel and sorted by nucleotide sequence. Sequence logos were assembled in R (version 2024.12.1+563, R Foundation for Statistical Computing, Vienna, Austria) using the ggseqlogo and Biostrings packages. Predicted VcircRNAs were also analyzed for trends in VcircRNA sequence length based on nucleotide calls and junction site breakpoints along the genome. To identify differential prediction hotspots or biases in genomic loci of predicted VcircRNAs by each algorithm, arc diagrams were produced using the SARS-CoV-2 reference genome (NCBI Accession #NC_045512.2) as the x-axis. Arc diagrams were assembled using ggplot2.

To quantify unique circRNAs predicted across algorithms, spreadsheets of unique VcircRNAs were compiled by detection method (algorithm) (Fig. S3A-B). Repeated detection of a unique VcircRNA by multiple algorithms was identified by sorting and counting intersections via dplyr (v. 1.1.4) and tidyr (v. 1.3.1) packages in R. Upset plots were generated using the UpSetR package (v. 1.4.0) in R. Overlapping predictions were more common for positive sense VcircRNAs (Fig. S3A) than in negative sense VcircRNAs (Fig. S3B), but overall, many of the algorithms identified unique putative VcircRNAs. The total number of unique VcircRNAs identified were used for these calculations rather than those above an arbitrary cutoff of read counts (as proxy for abundance), as many of the VcircRNAs predicted by multiple algorithms were predicted with different read counts across algorithms, suggesting variation in sensitivity. There were no VcircRNAs that were predicted by all four algorithms, regardless of strandedness or genome location. Find_circ and circRNA_finder had the highest number of overlapping predictions (202) for positive strand VcircRNAs. Only a cumulative 39 positive strand VcircRNAs and cumulative 13 negative strand VcircRNAs were predicted by three of the four algorithms. The largest number of unique VcircRNAs predicted was by SICILIAN, which is not designated for circRNA-specific predictions, but rather broader species produced by alternative splicing, and may indicate a larger number of false positives reported or less stringency with determining circRNA species.

### SARS-CoV-2 VcircRNA in acute and chronic clinical samples

Additional samples were acquired to investigate whether circ7b8N is detectable at different stages of SARS-CoV-2 infections (acute versus post-acute). Samples were acquired from individuals acutely infected with SARS-CoV-2 and longitudinally post-infection via nasopharyngeal (NP) swabs, fecal collection, and venipuncture for peripheral blood mononuclear cell (PBMC) collection. NP and fecal samples were collected as a part of the Lambda clinical trial conducted at Stanford University in 2020[23,25]. Sample collection and RNA extraction was conducted as previously described. PBMCs were acquired by the Stanford Long COVID Center, and RNA extraction was performed using the RiboPure RNA extraction kit from Ambion (#AM1924, Thermo Fisher Scientific, Fremont, CA, USA) following the manufacturer’s protocol. RNA eluents for all clinical samples were aliquoted and stored at −80 °C.

### Confirmation of circ7b8N

Primer design: Primers specific to the circ7b8N sequence were designed for PCR-based amplification. More specifically, a reverse (R) primer (CATGTTCGTTTAGGCGTGACA) was designed upstream of the 5’ breakpoint so that it would amplify products 3’ → 5’, in the direction of the 5’ end of the junction site. The R primer can anneal to either circular or linear RNA products and is used for downstream cDNA synthesis and PCR amplification. A forward (F) primer (TGATTACAAACTTCTGCTATTCCTTGT) was designed to span the junction site, which is unique to the circRNA permutation, and is not found contiguously in the linear sequence. Primers were synthesized by the Stanford University Protein and Nucleic Acid (PAN) Facility.

Ribonuclease R (RNase R) treatment: RNase R is an exoribonuclease enzyme to which many circRNAs are resistant, as it degrades RNA through hydrolysis in the 3’ → 5’ direction. RNase R (# RNR07250, LCG Biosearch Technologies, Novato, CA, USA) was used to treat RNA samples to reduce linear RNA amplification and enhance circRNA availability, while also confirming resistance to degradation via later amplification steps. A reaction mixture of 15ug of RNA was added to 2.5 μl of RNase R enzyme, 5.5 μl of RNase R reaction buffer, and sterile, nuclease-free water to a total volume of 10 μl. Reaction mixes were pipetted briefly to mix, centrifuged to concentrate the reaction mix to the bottom of the tube, and incubated in a thermocycler at 37 °C for 15 minutes, followed by a 4 °C hold. Following RNase R treatment, RNase R was removed from the RNA samples using the Zymo RNA Clean and Concentrator kit (#R1019, Zymo Research Corporation, Irvine, CA, USA), following the manufacturer’s protocol. Once cleaned, the entire concentrated aliquot (25 μl) of RNase R-treated, cleaned RNA was used for cDNA synthesis.

Rolling circle amplification for cDNA synthesis: Complementary DNA (cDNA) was synthesized from RNA extracted from infected A549-ACE2 cells using the R primer and the High-Capacity cDNA to RNA kit from Applied Biosystems (#43-874-06), Thermo Fisher Scientific, Waltham, MA, USA). Briefly, 1 μl of untreated RNA was added to a reaction mix of reverse transcriptase (RT) enzyme (0.5 μl), 2x reaction buffer (5 μl), 10 μM R primer (0.2 μl), and sterile, nuclease-free water (3.3 μl) for a total volume of 10 μl per reaction. If generating cDNA from RNase R-treated RNA, 25 μl of RNase R-treated and cleaned RNA was added to a reaction mix of RT enzyme (0.5 μl), 2x reaction buffer (5 μl), and 10 μM R primer (0.2 μl). Reaction mixes were mixed by pipetting and centrifuged briefly before incubating in a thermocycler at 37 °C for 60 minutes. Following incubation, the reaction was heated to 95 °C for 5 minutes, then held at 4 °C. cDNA samples were either frozen at −20 °C until further processing or transferred to ice and used immediately.

Targeted amplification of circ7b8N cDNA by polymerase chain reaction (PCR): PCR was used to amplify circRNA junction sites within the cDNA generated from SARS-CoV-2 infection samples. The CloneAmp HiFi PCR Premix from Takara (#639298, Takara Bio USA, San Jose, CA, USA) was used as a master mix. In brief, 12.5ml of HiFi PCR Premix was added to 1 μl cDNA, 1 μl each of circle-specific F and R primers, and 9.5 μl of nuclease-free water to bring the reaction volume to 25 μl. Master mix aliquots were vortexed briefly and centrifuged after aliquoting into PCR tubes, and again after adding cDNA. The PCR reaction was performed in a thermocycler, where samples were heated to 98 °C for 1 minute, followed by 35 rounds of denaturation (98 °C for 10 seconds), annealing (55.5 °C for 15 seconds), and extension (72 °C for 1 minute), a single extended 72 C hold for 5 minutes to complete a final extension, and a 4 °C indefinite hold until samples could be either moved to – 20 °C for storage or transferred to ice until processing by gel electrophoresis.

Gel electrophoresis for PCR visualization: Agarose gels (0.8% concentration) were made with 1X Tris-Borate-EDTA (TBE) buffer. TBE-agarose mixtures were heated in a microwave until agarose was completely melted and cooled to handling temperature before adding 5 μl of GelGreen Nucleic Acid Dye (#41005, Biotium, Inc., Fremont, CA, USA). Gels were set in a gel cast frame at room temperature until firm and run in a gel tank submerged in 1X TBE buffer. Gels were imaged using a ChemiDoc™ MP Imaging System (#12003154, BioRad, Hercules, CA, USA). Appropriate-sized bands were extracted from agarose gels using the QIAquick gel extraction kit (#28704, Qiagen US, Germantown, MD, USA) following the manufacturer’s protocol.

TOPO cloning for sequence confirmation of rolling circle amplification: DNA bands extracted from electrophoresis gels were each cloned into plasmids by topoisomerase (TOPO) cloning. Using the Ultra-Universal TOPO cloning kit with a pCE3 blunt vector, both from Vazyme (#C603, Vazyme Biotech, Nanjing, China), each band was ligated into the pCE3 vector backbone by adding the optimized concentration of insert (according to the Vazyme TOPO protocol, up to 8 μl volume) to 2 μl of 5X Ultra-Universal TOPO Cloning mix, and sterile, nuclease-free water to bring the final reaction volume to 10 μl. Reaction mixes were mixed by pipetting and centrifuged to concentrate the reaction to the bottom of the tube. The reaction was then performed by incubation in a thermocycler at 25 °C for 5 minutes. Reaction tubes were then placed on ice before being used to transform Stellar chemically competent cells (#636763, Takara Bio USA, Inc., San Jose, CA, USA). Stellar cells were thawed on ice and aliquoted to 50 μl per transformation tube. The entire 10 μl reaction was added to an aliquot of Stellar cells and mixed by gentle stirring with a pipette tip. Once mixed, the cells were incubated on ice for a minimum of 30 minutes. Tubes were then heat-shocked at 42 °C for 45 seconds and immediately placed on ice again for 2 minutes. Warmed SOC media was added to round bottom falcon tubes (440 μl per tube), and each competent cell reaction was added entirely (60 μl total), for a total of 500 μl per falcon tube. Tubes were incubated at 37 °C, shaking, for 60 minutes to allow for transformed cell population expansion. From each expansion, 100 μl of expanded cells was added to a LB media agar plate made with carbomycin antibiotic for plasmid selection, spread, and incubated overnight at 37 °C. The following day, distinct colonies were selected from each plate and added to 3 ml of LB liquid media with carbomycin for further selection and expansion by overnight shaking incubation at 37 °C. TOPO plasmids were extracted using the QIAGEN plasmid mini kit (#12123, QIAGEN, Germantown, MD, USA).

Whole plasmid sequencing: Plasmids extracted from TOPO cloning events were sent to Plasmidsaurus to confirm the success of the cloning by Whole Plasmid Sequencing (WPS) by Oxford Nanopore Technology with custom analysis and annotation (Plasmidsaurus, Eugene, OR, USA). The resulting GenBank files were analyzed using SnapGene software (v. 8.0+, GSL Biotech LLC, Boston, MA, USA).

### Synthesized circRNA and expression vectors for circ7b8N overexpression

An expression vector of circ7b8N was generated using Gibson cloning techniques. The pcDNA3.1(+) vector backbone contains two laccase2 inverted repeats to stimulate circularization of the DNA insert[69]. The pcDNA3.1(+) Laccase2 MCS Exon Vector was a gift from Jeremy Wilusz (Addgene plasmid # 69893; http://n2t.net/addgene:69893; RRID: Addgene_69893). Insert sequences were synthesized by Twist Biosciences (Twist Biosciences, South San Francisco, CA, USA). To ensure uninterrupted detection of the natural junction site, the circ7b8N sequence was split at a site 208 nucleotides into ORF N to generate the blunt ends required for cloning. We chose to insert the breakpoint within the ORF to also preserve the transcription regulatory sequences (TRSs), which flank ORF 8 in circ7b8N, in case they conferred any significant function.

Synthesized circRNA of circ7b8N was produced by GenScript (GenScript USA, Inc., Redmond, WA, USA) using the same split sequence as the insert described above, wherein the natural junction site and TRSs are uninterrupted. Synthesized circ7b8N was purified by high performance liquid chromatography (HPLC) by GenScript prior to delivery.

The synthesized circ7b8N from GenScript and the circ7b8N generated by the circ7b8N expression vector were both confirmed to be detectable by the existing R and F primer set, resistant to RNase R treatment, and confirmed by sequencing to ensure matching RNA sequences to the originally predicted and confirmed circ7b8N sequence. The results reported in this paper all reflect transfections with the GenScript synthesized circ7b8N.

### Viral kinetics

Viral titering and quantitative PCR (qPCR) were performed to assess whether the presence of circ7b8N influences viral replication. A549-hACE2 cells were seeded into 12-well plates and grown to ∼80% confluency overnight. Then, each well was transfected with synthesized circ7b8N in 2.0 μg, 1.0 μg, 0.5 μg, and 0.25 μg concentrations in triplicate using the X-fect RNA Transfection Reagent from Takara (#631450, Takara Bio USA, Inc., San Jose, CA, USA) and following the manufacturer’s protocol. Transfected cells were incubated in the transfection media at 37 °C for 4 hours before replacing the transfection media with complete DMEM growth media. Once media was replaced, transfected cells were incubated overnight at 37 °C with 5% CO_2_. Cells were transferred to the BSL-3 facility for infection at 24-hours post transfection. Cells were infected with SARS-CoV-2 isolate USA-WA1/2020 at an MOI of 0.5. After 1 hour at 37 °C, infection inoculum was removed, cells were rinsed with PBS and overlaid with new DMEM growth media. After 24 HPI (and 48-hours post transfection of circ7b8N), supernatants were collected and titrated by plaque assay as described previously[70]. TRIzol Reagent (#15596026, Thermo Fisher Scientific, Waltham, MA, USA) was added to the cells and scraped into tubes for RNA extraction. Downstream RNA extraction was completed using the Direct-Zol RNA miniprep kit from Zymo Research (#R2052, Zymo Research Corporation, Irvine, CA, USA).

Viral RNA isolated from each transfection condition was quantified by quantitative PCR (qPCR). Prior to qPCR, 100 ng of each RNA sample was used as a template for cDNA generation with a target-specific R primer, as described earlier. To quantify viral RNA as an indicator for viral replication, instead of using primers to target circ7b8N, the RNA-dependent RNA polymerase (RdRp), encoded in ORF12, was amplified using a target-specific R primer (5’-GCCGTGACAGCTTGACAAAT-3’). A GAPDH-specific R primer (R: 3’-GGGGAGATTCAGTGTGGTGG-5’) was used to amplify GAPDH as a housekeeping gene. Taq Pro Universal SYBR qPCR Master Mix from Vazyme (#Q712-02, Vazyme Biotech, Nanjing, China). The qPCR reaction mix included 10 ml of 2x Taq Pro Universal SYBR qPCR Master Mix, 2 ml of cDNA, and 7.2 ml of sterile, nuclease-free water. Primer pairs for RdRp (F: 5’-GTGARATGGTCATGTGTGGCGG-3’, R: 5’-GCCGTGACAGCTTGACAAAT-3’) or GAPDH (F: 5’-TTGGCTACAGCAACAGGGTG-3’, R: 3’-GGGGAGATTCAGTGTGGTGG-5’) were added to each master mix at 0.4 μl from 10 mM stocks. The qPCR reaction included an initial denaturation stage of a 95°C incubation for 30 seconds, followed by 40 repetitions of a cycling reaction stage (95°C for 10 seconds, 60°C for 30 seconds), and a melting curve stage (95°C for 15 seconds, 60°C for 1 minute, and 95°C for 15 seconds). Cq values were analyzed for target expression.

### Cellular localization of circ7b8N by Fluorescent In-Situ Hybridization (FISH)

FISH was employed using the ThermoFisher Scientific ViewRNA Cell Plus Assay kit (#88-19000-99, ThermoFisher Scientific, Waltham, MA, USA) combined with a specialized probe targeting the junction site of circ7b8N. Probes were tagged with an Alexa Fluor 546 and designed to straddle the junction site (ACUGAUUACAAACAnnnnnUUUCUGCUAUUCCUU). In short, A549-hACE2 cells were seeded in the wells of a 24-well plate on a sterile coverslip in complete DMEM with 50 mg/ml Geneticin (#10131035, ThermoFisher Scientific, Waltham, MA, USA) and grown overnight to ∼80% confluency. The following day, cells were introduced to synthesized circ7b8N or SARS-CoV-2 WA.1 strain live virus. Untransfected and uninfected A549-hACE2 cells were used as a negative control. An additional control of a synthesized eGFP encoding circRNA with the Coxsackie B virus internal ribosomal entry site (IRES) was used as a specificity test for the FISH probe designed against the circ7b8N junction site. For synthesized circ7b8N and replicon IVT RNA: Once confluent, growth medium was aspirated, and cells were washed briefly with 1x PBS. After washing, 500 μl of Opti-MEM media was added to each well. Cells were transfected with synthesized circ7b8N or IVT RNA using the Takara Xfect RNA transfection kit (#631450, Takara Bio USA, Inc., San Jose, CA, USA) following the manufacturer’s protocol for 24-well transfections. For SARS-CoV-2 live infection: Once confluent, growth medium was aspirated, and cells were washed with 1X PBS. MOI was calculated (0.5 MOI), and SARS-CoV-2 isolate USA-WA1/2020 (#NR-52281, Biodefense and Emerging Infections (BEI) Resources, Newport News, VA, USA), initially grown to passage 3 in VeroE6-TMPRSS2 cells and isolated, was added to A549-hACE2 cells. Once the virus was added, cells were incubated at 37 °C for one hour to allow for viral attachment. Following incubation, inoculum was removed by aspiration and cells were rinsed with 1X PBS and fresh complete DMEM was added to each well. Cells were incubated at 37 °C until harvesting (72 HPI). The ViewRNA Cell Plus Assay procedures were performed over three days as follows:

Day 1: Fixation and Permeabilization: Growth medium was aspirated, and cells were incubated in Fixation/Permeabilization Solution for 30 minutes at room temperature. Cells were then thrice washed in 1X PBS with RNase Inhibitor. Once washed, cells were stored in 1X PBS with RNase Inhibitor at 4 °C overnight.

Day 2: Target Probe Hybridization: PBS was aspirated from cells and replaced with ViewRNA Cell Plus Probe Solution containing the circ7b8N custom probe. Once immersed in probe/probe solution, cells were incubated at 40 °C in a humidified incubator for 2 hours. Following incubation, probe solution was aspirated, and cells were washed in ViewRNA Cell Plus RNA Wash Buffer Solution five times. Once washed, cells were stored in ViewRNA Cell Plus RNA Wash Buffer Solution at 4 °C overnight.

Day 3: Signal Amplification: Samples were removed from overnight incubation and warmed to room temperature. While warming, ViewRNA Cell Plus Amplifier Diluent and ViewRNA Cell Plus Amplifier Mix were warmed to 40 °C for ∼30 minutes. ViewRNA Cell Plus Pre-Amplifier Mix, Amplifier Mix, and Label Probe Mix were thawed to room temperature and stored on ice until ready to use. Wash buffer was aspirated from each sample well and ViewRNA Cell Plus Pre-Amplifier Solution was mixed to a 1:25 dilution and added to each well. Samples were incubated in Pre-Amplifier solution for 1 hour at 40 °C in a humidified incubator. Following incubation, samples were washed five times with ViewRNA Cell Plus Wash Buffer Solution. Once washed, the ViewRNA Cell Plus RNA Amplifier Mix was prepared to a 1:25 dilution and added to each sample. Samples were incubated in Amplifier solution for 1 hour at 40 °C in a humidified incubator. Following incubation, samples were washed five times with ViewRNA Cell Plus Wash Buffer Solution. Following wash steps, the ViewRNA Cell Plus Working Label Probe Mix Solution was prepared to a 1:25 dilution and added to each sample well. Samples were incubated in Working Label Probe Mix Solution for 1 hour at 40 °C in a humidified incubator. Following incubation, samples were washed five times with ViewRNA Cell Plus Wash Buffer Solution and allowed to incubate at room temperature in the final wash for 10 minutes prior to final aspiration. Once washed, samples were gently washed in 1X PBS and stained with DAPI and phalloidin (1:100 dilution for each in 1x PBS) for 10 minutes. After DAPI and phalloidin staining, cells were washed again in 1X PBS. To mount each sample, glass slides were prepared with mounting media and glass coverslips with fixed cells adhered were placed cell-side down on the glass slide. Each coverslip was sealed around the edges with clear nail polish, and slides were stored in the dark at 4 °C until analysis. Slides were imaged using 63x lens and channels on a Zeiss LSM 700 laser scanning confocal microscope. Images were processed using Fiji (https://imagej.net/software/fiji/).

### Circ7b8N influence on host cellular responses

To establish whether circ7b8N alters host or SARS-CoV-2 gene expression, A549-hACE2 cells were exposed to multiple conditions in triplicate: 1) circ7b8N transfections only, which were transfected with 2 μg of circ7b8N for 24 hours before RNA was extracted, 2) infection only, where cells were infected with SARS-CoV-2 WA1/2020 for 24 hours before RNA extraction, 3) “overexpression”, wherein cells were transfected with circ7b8N directly for 24 hours, followed by infection with SARS-CoV-2 WA1/2020 for 24 hours before RNA extraction, and 4) uninfected and untransfected cells as a negative control. All transfections were performed with the Xfect RNA transfection reagent from Takara () as previously described. All infections were performed as previously described. RNA from each condition was sent to Genewiz (Genewiz from Azenta Life Sciences, South Plainfield, NJ, USA) for library preparation and bulk RNAseq using Next Generation Sequencing (NGS). NGS RNAseq was performed with rRNA depletion and without poly-A selection.

Raw FASTA files from each sequenced sample were trimmed using Trim Galore! (v. 0.5.0, https://www.bioinformatics.babraham.ac.uk/projects/trim_galore/) to remove Illumina adapter sequences, followed by quality control assessment using FastQC (https://www.bioinformatics.babraham.ac.uk/projects/fastqc/). Once the quality of the reads was verified, each trimmed FASTA file was aligned to an indexed human reference genome, <u>GRCh38.p14</u>. Genome sequence and comprehensive gene annotation files were acquired from GenCode (European Molecular Biology Laboratory - European Bioinformatics Institute (EMBL-EBI), and were indexed prior to alignment using STAR. Aligned reads were quantified by RNA-Seq by Expection-Maximization (RSEM)[71] (v. 1.3.3, https://deweylab.github.io/RSEM/). RSEM outputs were used as inputs for DESeq2[72] to analyze differential gene expression across conditions. DESeq2 results were visualized by volcano plots generated by ggplot2[73] (https://ggplot2.tidyverse.org/), and the resulting lists of differentially expressed genes were sorted by False Discovery Rate (FDR)[74]. Genes and pseudogenes with an adjusted p-value of ⋜0.05 or ⋜0.01 were ranked by significance and submitted to gProfiler[75] (https://biit.cs.ut.ee/gprofiler/gost, version *e113_eg59_p19_f6a03c19*, database updated on 23/05/2025) for an ordered query against the *Homo sapiens* genome. Enriched GO term results from gProfiler queries were trimmed of redundant terms and term relationships (i.e. significantly upregulated or downregulated) were visualized using ggplot2[73] (https://ggplot2.tidyverse.org/).

### Circ7b8N influence on viral gene expression

Differential viral gene expression in experimental conditions was assessed by computational analysis of the RNAseq data described above. FASTA files generated by NGS RNAseq were trimmed and subjected to FastQC as previously described. The SARS-CoV-2 reference genome and annotation files were acquitted from NCBI (Accession #<u>GCF_009858895.2)</u> and indexed for alignment with Salmon[76] (https://combine-lab.github.io/salmon/). Trimmed RNAseq reads were aligned and viral gene expression was quantified with Salmon across all experimental conditions. Salmon quantified output data was transformed for visualization using tximport[77] (DOI: <u>10.18129/B9.bioc.tximport</u>) and DESeq2[72], and bar plot was generated using ggplot2[73] (https://ggplot2.tidyverse.org/).

### Assessment of circ7b8N conservation

Clustal-Omega multisequence alignments were performed using the Sequence Analysis Tools Job Dispatcher from EMBL European Bioinformatics Institute (EMBL-EBI), and results were analyzed using JalView (v. 14.6.1). Downstream analyses of point mutations were completed in SnapGene (v. 8.0.3). Needle pairwise alignments of protein sequences for NSP1 were conducted using the EMBL-EBI Job Dispatcher.

To assess circ7b8N generation in early SARS-CoV-2 variants, cultures of Caco-2 (human colon adenocarcinoma) and Calu-3 (human lung adenocarcinoma) cell lines were infected with variant isolates of SARS-CoV-2 and ORF deletion mutant strains (dORF) of the WA.1 SARS-CoV-2 strain, as previously described. Briefly, SARS-CoV-2 strains (WA-1 strain, Alpha, Delta, Omicron BA.1 and BA.5) were obtained from BEI Resources[78,79]. SARS-CoV-2 deletion mutants were kindly provided by Dr. Matthew B Frieman at University of Maryland[80,81]. Virus stocks were prepared by infecting VERO-TMPRSS2 cells in media containing 2% FBS supplemented with HEPES[82]. At 3 dpi, the cells were subjected to freeze-thawed, and media was collected and centrifuged 3,000 rpm for 20 min. The supernatant containing virus was aliquoted and stored at −80°C as the seed stock (P0). The P0 stock was sequence-verified, amplified in VERO E6 cells to generate working stock (P1), and used for all experiments. The titer of virus stock was determined by a 50% tissue culture infective doses (TCID50) using the Reed-Muench method in VERO E6 cells as described previously[82]. All work with live SARS-CoV-2 was performed in a biosafety level 3 (BSL3) laboratory and approved by the Institutional Biosafety Committee and Environmental Health and Safety at University of Pennsylvania.

Calu-3 cells were plated onto collagen-coated plates and inoculated with SARS-CoV-2 at a multiplicity of infection (MOI) of 0.1 one day post-plating. At 2 days post-infection (dpi), the cells were harvested using TRIzol reagent (Invitrogen), and total RNA was purified using the RNA Clean & Concentrator-25 kit (Zymo Research). Similarly, Caco-2 cells were seeded onto collagen-coated plates and inoculated with SARS-CoV-2 (MOI 0.1) three days post-seeding. Total RNA from the Caco-2 cells was extracted at 2 dpi following the identical protocol used for the Calu-3 cells.

RNA isolates were received as a gift and were processed by rolling circle amplification for cDNA generation with circ7b8N-specific R primer, both with and without prior RNase R treatment, followed by PCR amplification with circ7b8N-specific F and R primers. PCR products were visualized by gel electrophoresis, and bands coinciding with circ7b8N size (1372 nucleotides) were excised for DNA extraction. DNA extraction was performed with the QIAquick gel extraction kit (#28704, Qiagen US, Germantown, MD, USA) following the manufacturer’s protocol. Extracted DNA products were TOPO cloned and sent for WPS to confirm the presence of circ7b8N in Caco-2 and Calu-3 cell infections with different SARS-CoV-2 variants.

Reference genomes for the ancestral strain (“WuHu-1 2019”, Accession #NC_045512), ancestral isolate (WA.1 2020, Accession #MT246667.1), and Omicron (Omicron BA.1, Accession #ON026021) were accessed from NCBI. Representative genomes for clades Alpha (EPI ISL #601443), Beta (EPI ISL #712073), and Delta (EPI ISL#3148365) were accessed through GISAID EpiCoV Database following the GISAID EpiCoV Database Access Agreement and were converted to RNA sequences using Geneious Prime (Geneious Prime release 2024.0.5). Whole sequences for WA.1, Alpha, Beta, Delta, and Omicron BA.1 were aligned to the WuHu-1 reference genome using Geneious Prime assembler function. An extraction of the circ7b8N region, plus the 14 nucleotides which flank the 5’ and 3’ junction site shows that the predicted junction region is conserved across all 6 assembled sequences. While reference genomes represent a snapshot of the viral sequence within very specific conditions, the conservation of the junction site further supports our detection of circ7b8N across many variants and clinical samples and suggests a possible significance with VcircRNA for SARS-CoV-2 pathogenesis.

### Ethics Statement

The trial was registered at ClinicalTrials.gov (NCT04331899). The study was performed as an investigator initiated clinical trial with the FDA (IND 419217). It was approved by the Institutional Review Board of Stanford University (IRB) protocols # 55619 (PIs: Upinder Singh, Prasanna Jagannathan) and # 42043 (PI: Ami Bhatt). The data utilized has been stripped of personal identifiers.

## Supplemental Figures

**Figure S1: Arc diagrams and junction motifs of CIRI2, circRNA_finder, and find_circ algorithm predictions.** Positive (maroon) and negative (light blue) arcs indicate circularized regions of the SARS-CoV-2 genome, as predicted by CIRI2 (A), circRNA_finder (C), and find_circ (E). Grey dots at the end of each arc indicate breakpoint positions along the viral genome. Sequence logos were constructed for each algorithm used to predict VcircRNAs from SARS-CoV-2 RNAseq data. Probabilities were calculated for each nucleotide within 10 positions from the terminal end of each breakpoint for predicted positive and negative sense circRNAs separately to identify any unique, stranded motifs. (B) CIRI2 favors Guanine (G) in many of the terminal-most nucleotides in both 3’ and 5’ breakpoints, except in negative sense 5’ breakpoints. (D) CircRNA_finder did not have a favored nucleotide in the terminal nucleotides for each breakpoint. Negative sense circRNA prediction motifs for circRNA_finder were limited to n=2 predicted circRNAs and do not hold statistical weight but are only shown for a visual comparison. (F) Find_circ also did not present any prominent motifs, except with a slight favorability towards Uracil (shown as T) in positions flanking the immediate junction site breakpoint in the positive circRNA predictions.

**Figure S2: Numerical comparison of unique VcircRNAs predicted by different algorithms.** Upset plots of (A) positive sense and (B) negative sense VcircRNAs predicted by SICILIAN, CIRI2, circRNA_finder, and find_circ algorithms. Black dots indicate representations along the top bars, illustrating the number of unique circRNAs predicted exclusively by an algorithm. Black dots connected by black bars represent the number of unique circRNAs predicted across multiple algorithms. Grey dots indicate a particular algorithm did not contain overlapping predictions with other algorithms in each category and is therefore not represented in that cluster.

**Figure S3: Predicted secondary structure of circ7b8N.** Produced using RNAfold WebServer.

**Figure S4: Plasmid maps confirming circ7b8N expression during acute infection.** RNA extracted from NP swabs collected during acute infection was analyzed by rolling circle amplification, PCR, and gel electrophoresis for circ7b8N expression (Fig. 3A). Bands suggesting the presence of circ7b8N were extracted, TOPO cloned and sequenced by whole plasmid sequencing. Each plasmid map confirms the presence of circ7b8N by the presence of two junction sites (neon green) flanking the entire circRNA sequence, with two copies of 7b (olive green), ORF8 (yellow), and N fragment (light blue) inserted into the TOPO vector sequence (grey).

**Figure S5: Viral replication in response to circ7b8N treatment.** (A) A549-hACE2 cells were treated with circ7b8N for 12 hours by RNA transfection. At 12 hours, cells were infected with SARS-CoV-2. Supernatant was collected at 15 hpi and used for titering by plaque forming assay. Treatment with circ7b8N at any concentration did not significantly alter viral titer. (B) Quantification of viral E protein in response to treatment with circ7b8N prior to infection was assessed by qPCR. Statistical analyses performed using a one-way ANOVA and pairwise t-tests against the virus-only condition.

**Figure S6: Plasmid maps confirming circ7b8N expression by SARS-CoV-2 variants in Calu3 and Caco2 cells.** RNA extracted from Calu3 and Caco2 cell lines infected with variants of SARS-CoV-2, as well as WA.1 mutants dORF3a, dORF6, dORF7a, and dORF8, was analyzed by rolling circle amplification, PCR, and gel electrophoresis for circ7b8N expression (Fig. 7). Bands suggesting the presence of circ7b8N were extracted, TOPO cloned and sequenced by whole plasmid sequencing. Each plasmid map confirms the presence of circ7b8N by the presence of two junction sites (neon green) flanking the entire circRNA sequence, with two copies of 7b (olive green), ORF8 (yellow), and N fragment (light blue) inserted into the TOPO vector sequence (grey), with the exception of WA.1 dORF8, which contained a truncated version of circ7b8N without ORF8 illustrated by two copies of the junction site (neon green) with two copies of 7b (olive green) flanking the N fragment (light blue) in the TOPO vector sequence (grey).

**Figure S7: Conservation of circ7b8N across *Coronaviridae* genera.** (A) MSA of CoVs of junction site breakpoints for circ7b8N. Sequences are sorted along the y-axis by viral genus (Deltacoronaviruses, Alphacoronaviruses, Gammacoronaviruses, and Betacoronaviruses). Two sequences (Bat CoV/PREDICT/PDF-2180 and Bat CoV) lacked assignment into genus categories, but they are likely of the Betacoronavirus genus. Consensus sequences below each alignment composite indicate no point mutations within the junction site (red arrows), yet minimal coverage across genera. Red arrows at the bottom of the alignment indicate the direction of the circRNA sequence from each breakpoint. The 2019 Hu.1 strain (highlighted in black, bottom) was used as the reference for each MSA. Library preparations for RNAseq convert Uracil (U) to Thymine (T), as reported here. (B) Phylogenetic analysis of genomes used to assemble the pan-coronovirus MSA in panel A. Genomes labeled in teal contain circ7b8N.

